# Essential strategies for the detection of constitutive and ligand-dependent Gi-directed activity of 7TM receptors using bioluminescence resonance energy transfer

**DOI:** 10.1101/2024.12.04.626681

**Authors:** Sofia Endzhievskaya, Kirti Chahal, Julie Resnick, Ekta Khare, Suchismita Roy, Tracy M. Handel, Irina Kufareva

## Abstract

The constitutive (ligand-independent) signaling of G protein-coupled receptors (GPCRs) is being increasingly appreciated as an integral aspect of their function; however, it can be technically hard to detect for poorly characterized, e.g. orphan, receptors of the cAMP-inhibitory Gi-coupled (GiPCR) family. In this study, we delineate the optimal strategies for the detection of such activity across several GiPCRs in two cell lines. As our study examples, we chose two canonical GiPCRs - the constitutively active Smoothened and the ligand-activated CXCR4,-and one atypical GPCRs, the chemokine receptor ACKR3. We verified the applicability of three Bioluminescence Resonance Energy Transfer (BRET)-based assays - one measuring changes in intracellular cAMP, another in Gβγ/GRK3ct association and third in Gαi-Gβγ dissociation, - for assessing both constitutive and ligand-modulated activity of these receptors. We also revealed the possible caveats and sources of false positives, and proposed optimization strategies. All three types of assays confirmed the ligand-dependent activity of CXCR4, the controversial G protein incompetence of ACKR3, the constitutive Gi-directed activity of SMO, and its modulation by PTCH1. We also demonstrated that PTCH1 promotes SMO localization to the cell surface, thus enhancing its responsiveness not only to agonists but also to antagonists, which is a novel mechanism of regulation of a Class F GiPCR Smoothened.

## Introduction

Due to their crucial roles in mammalian cell signaling and the accessibility of their ligand-binding pockets, members of the G protein-coupled receptor (GPCR) family form the center of the druggable proteome [1,2]. GPCR signaling is triggered by diverse types of signaling molecules (agonists); additionally, more than 60 GPCRs [3] have been reported to exert agonist-independent or *constitutive* (*basal*) activity. Excessive GPCR basal signaling, through gene amplification or gain-of-function mutations, is implicated in various human disorders [4–6]. Therefore experimental detection and quantification of such activity is critical for understanding GPCR pharmacology and for drug discovery.

GPCRs initiate intracellular signaling by coupling to heterotrimeric G proteins, composed of an obligate Gβγ heterodimer [7,8] and a Gα subunit of one of four subtypes: cAMP-stimulatory (Gαs), cAMP-inhibitory (Gαi/o), phospholipase C-activating (Gαq/11) or Rho GTPase-activating (Gα12/13) [9,10]. cAMP-inhibitory Gαi/o proteins are believed to mediate signaling of at least 40% of GPCRs [11,12]. In the inactive state, a Gα subunit is bound to guanosine diphosphate (GDP) which allows it to form a stable trimer with Gβγ (**Fig. 1A)**. Activated GPCR triggers a conformational change in the Gα subunit [13–15] that results in it releasing GDP and taking up guanosine triphosphate (GTP) that is about 10 times more abundant in the cytosol [16]. Because GTP-bound Gα can no longer form a trimer with Gβγ [17], this leads to the dissociation of Gα-GTP from the Gβγ heterodimer and subsequent modulation of downstream effectors by both Gα-GTP and Gβγ [7,8] (**Fig. 1A)**. Adenylyl cyclases (ACs) are downstream effectors of GTP-bound Gα subunits of both Gs and Gi proteins, resulting in AC activation and inhibition respectively, with subsequent elevation or suppression of cellular cAMP levels [7,8] (**Fig. 1A)**. Concurrently, Gα-free Gβγ heterodimers bind and engage numerous effectors of their own, one class of such effectors being the GPCR kinases (GRKs) 2 and 3 which directly bind Gβγ via their C-terminal pleckstrin homology (PH) domains [18–21].

**Figure 1.**
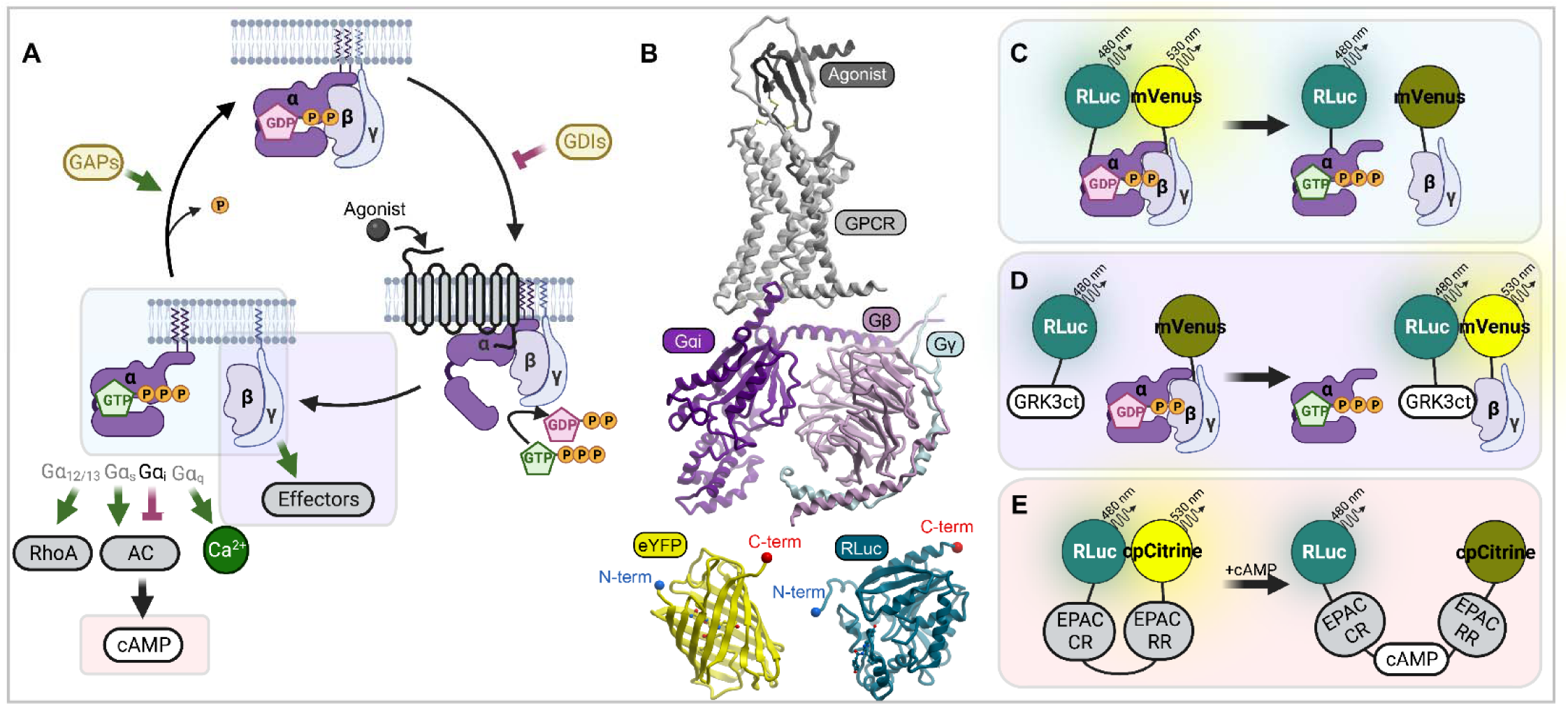
Monitoring the activation of Gi proteins downstream of GPCRs using Bioluminescence Resonance Energy Transfer (BRET). (**A**) Schematic representation of activation-signaling-deactivation cycle of heterotrimeric G-proteins. (**B**) AlphaFold2 model of a GPCR (CXCR4) with its agonist (CXCL12) in complex with Gαi, Gβ and Gγ next to crystal structures of eYFP and RLuc. Proteins are shown to scale. (**C**) Principle of a BRET-based assay measuring Gαi-Gβγ dissociation. (**D**) Principle of a BRET-based assay measuring the association between Gβγ released from activated Gαi, and the Gβγ effector GRK3. (**E**) Principle of an assay that utilizes a unimolecular BRET-based biosensor to measure the concentration of intracellular cAMP. AC, adenylyl cyclase. GTP, Guanosine Triphosphate. GDP, Guanosine Diphosphate. GDI, Guanine Nucleotide Dissociation Inhibitor. GAP, GTPase-Activating Protein. RhoA, Ras Homolog Family Member A. RLuc, Renilla Luciferase. GRK3ct, G Protein-Coupled Receptor Kinase 3 C-Terminus. EPAC CR, Exchange Protein Directly Activated by cAMP (Catalytically Reduced). EPAC RR: Exchange Protein Directly Activated by cAMP (Regulatory Reduced). cAMP, Cyclic Adenosine Monophosphate. cpCitrine, Circularly Permuted Citrine. mVenus, Monomeric Venus.

Capitalizing on these signaling pathways, various assays have been developed for studying ligand-induced GPCR signaling to their effector G proteins (e.g. GTPγ binding and various second messengers assays) [22]. A widely used class of assays is based on measuring inter-and intramolecular proximities in live cells using Bioluminescence Resonance Energy Transfer (BRET) [23–26]. The advantages of BRET assays include their adaptability to high throughput formats, the ability to monitor rapid changes in real time (vs end-point measurements), and the ratiometric (vs intensiometric) nature of the readout that is minimally affected by variations in reporter expression levels. However, because BRET ratios are always relative and hard to calibrate against any absolute quantity (e.g. the intracellular concentration of Gα/Gβγ heterotrimers), their use for characterization of GPCR *constitutive* activity is nontrivial and reliant on the availability of adequate comparisons and controls. The often low level of constitutive activity and its dependence on receptor expression and cell types additionally complicate the matter.

Numerous BRET-based G protein-directed activity reporters have been developed over the years, exploiting the different intracellular molecular interactions resulting from receptor activation (**Fig. 1A**). In terms of “proximity” to the coupling of the agonist-activated GPCR and G protein, the best BRET measurement is arguably between the receptor and the G protein itself; however, this assay requires receptor tagging with either a BRET donor or BRET acceptor, both relatively large (18-36 kDa, **Fig. 1B**) proteins with the potential to alter receptor signaling and/or trafficking [27,28]. Consequently, even though multiple GPCR-G protein BRET pairs have been developed and reported for well-characterized GPCRs [29–37], this design is less preferred for novel, orphan, and/or poorly characterized receptors [38,39].

Because of this, considerable efforts have been made to develop BRET reporters of G protein signaling that are suitable for use with untagged receptors. For such reporters, the most receptor-proximal event is the dissociation of Gα from Gβγ that can be measured as a decrease in BRET between donor-tagged Gα and acceptor-tagged Gβγ [40–43] (**Fig. 1C**). An advantage of this approach is being the most direct and thus unaffected by the potential interference from multiple integrated downstream pathways. One step away from the Gα-Gβγ dissociation event is a reporter where the acceptor is on Gβγ and the donor is on a Gβγ effector, or a fragment of thereof, such as the PH-domain containing C-terminus (ct) of GRK3 [44–48]: the release of Gβγ from Gα after GPCR activation allows Gβγ to interact with GRK3ct which results in elevated BRET (**Fig. 1D**). The advantage of this approach is that it can work with untagged WT or mutant Gα. Reporters that similarly detect GTP-bound (untagged or even endogenous) Gα have been developed as well [49–53]. Finally, the activation of two types of G proteins, Gs and Gi, alters the levels of intracellular cAMP and can be monitored with a BRET-based “**<u>cAM</u>**P sensor using the **<u>Y</u>**FP-**<u>E</u>**PAC-R**<u>L</u>**uc” (CAMYEL) reporter [54,55] or variations thereof [56,57] (**Fig. 1E**). In the **<u>E</u>**xchange **<u>P</u>**rotein **<u>A</u>**ctivated by **<u>c</u>**AMP 1 (EPAC 1, also known as RAPGEF3), cAMP-dependent activation changes the distance between the conserved N-terminal regulatory region (RR) and the C-terminal catalytic region (CR): in the absence of cAMP, the CR and RR are close but once cellular cAMP levels rise, they separate [55,58] (**Fig. 1E**). Capitalizing on this natural cAMP-dependent, conformation switching molecule, the CAMYEL cAMP reporter has an advantage of having both donor and acceptor in the same protein molecule (i.e. it is a unimolecular sensor), which eliminates variability associated with the levels of BRET acceptor and donor expression. CAMYEL and other unimolecular BRET cAMP biosensors are superior to luminescence-based alternatives such as GloSensor [59,60] because of their ratiometric nature, or CRE-Luc (cAMP response element coupled with a luciferase reporter [59,61,62]) because of their ability to track cAMP changes in real time.

Together with other cAMP biosensors, CAMYEL has been frequently used for characterizing the activity of Gs protein-coupled receptors (GsPCRs) whose activation leads to an *increase* in intracellular cAMP [63–65]. However, its application is less straightforward for Gi protein-coupled receptors (GiPCRs) whose activation leads to a *suppression* of intracellular cAMP [63,66–68], especially when such suppression is constitutive rather than agonist-induced. Additionally, because cAMP changes in the cell are at least two steps removed from the receptor-G protein coupling event, CAMYEL results can be affected by multiple integrated pathways and are often hard to deconvolute.

Head-to-head comparisons between these approaches and their ability to detect constitutive signaling of GiPCRs by monitoring events at varying “proximity” to the receptor activation event and in different cell lines have not yet been performed. In the following sections, we fill this knowledge gap and demonstrate the use and optimization of these assays for the detection of constitutive and ligand-dependent Gi-directed activity of several receptors in two cell lines. We identify important caveats, delineate cell-line specific optimization approaches, and suggest comparison strategies for receptors with limited ligands. Finally, with the use of the most “proximal” assay, Gαi-Gβγ dissociation, we reveal a novel regulatory mechanism for Smoothened (SMO), one of the receptors in our panel.

## Results

### Rationale for the present study

As a prototypical non-constitutively-active GiPCR-ligand pair, we chose the CXC chemokine receptor 4 (CXCR4) and its endogenous chemokine agonist CXCL12 (also known as SDF1). To dissect the various manifestations of constitutive Gi-directed activity, we included Smoothened (SMO), a cryptic Family F receptor and the central component of the Hedgehog signaling pathway that has been reported to be constitutively active towards Gi [60,62,69]. Its constitutive activity is suppressed by the 12-transmembrane (12TM) protein PTCH1 [60,70,71] and numerous small molecule antagonists including the approved antineoplastic (basal cell carcinoma) drug Vismodegib [72,73]. Finally, in selected assays we include the controversial atypical chemokine receptor 3 (ACKR3): despite sharing an agonist (CXCL12) and a high degree of sequence and structural homology to CXCR4, this receptor has been shown to signal through β-arrestin and not through G proteins [74–77]. However, conflicting reports regarding constitutive [78] or ligand-dependent [79,80] Gi-directed signaling have been published, prompting us to re-evaluate this receptor in our panel of assays using different Gi-directed activity reporters in two distinct cell lines. One of the cell lines that we chose to use is HEK293T, because of its widespread use in GPCR signaling literature [81–85] due to ease of maintenance and transfection. The related HEK293 cells have also been reported to express measurable levels of SMO and PTCH1 (Human Protein Atlas [86], accessed 2023-05-01, **Supp. Fig. 1A**) and hence may provide a suitable transcriptional environment for studies of SMO signaling. As an alternative, we selected HeLa, because this epithelial cell line endogenously expresses both CXCR4 and ACKR3 [76,87–91] (**Supp. Fig. 1B**) and migrates towards CXCL12 [92], thus providing an appropriate cellular context for these two receptors.

### A BRET-based cAMP reporter CAMYEL robustly detects constitutive and ligand-dependent activity of GiPCRs

A conventional BRET assay is conducted in a multi-well plate with live cells co-expressing proteins of interest, one tagged with a BRET donor (a luciferase such as RLuc) and another tagged with a BRET acceptor (e.g. YFP or mVenus). Luminescence is initiated by the addition of a compatible luciferase substrate (e.g. coelenterazine-h) after which BRET is measured as a ratio of emission in one part of the luciferase spectrum (e.g. yellow) to its emission in another part (e.g. blue or green) and monitored over time, in real-time, upon the addition of receptor stimuli or inhibitors. The increase in longer-to-shorter-wavelength emission ratio indicates resonant energy transfer that signifies that the donor and the acceptor get physically closer in the cell. In the CAMYEL assay because of the design of this unimolecular biosensor (**Fig. 1E**), it is more advantageous to monitor the inverse BRET, i.e. the shorter-to-longer-wavelength emission ratio, because this ratio changes monotonically (and not inversely) with the intracellular concentrations of cAMP.

The implementation of this assay in HEK293T cells co-expressing the CAMYEL cAMP sensor together with CXCR4, SMO, or with no exogenously transfected GPCR produces inverse BRET ratios that are indistinguishable between receptors (**Fig. 2A**) and are not affected by the addition of CXCR4 agonist CXCL12 or SMO antagonist Vismodegib (**Fig. 2B-D, second segment**). This is in striking contrast to CAMYEL-based studies of Gs-coupled receptors where the addition of receptor agonist immediately leads to a measurable elevation of cAMP and a change in the CAMYEL inverse BRET signal [63–65]. The basal CAMYEL inverse BRET ratio was also unaffected by pretreating cells with pertussis toxin (PTX, **Fig. 2A**), a naturally occurring toxin and a commonly used laboratory agent that, by ADP-ribosylating a Cys residue close to the C-terminus of Gαi (C351), abrogates the ability of cellular Gi proteins to couple to GPCRs [93–95]. However, notable changes in inverse BRET ratio are observed after the addition of forskolin (FSK) (**Fig. 2B**), a known AC activator and booster of intracellular cAMP. These changes are also indicative of the activation of the transfected GiPCRs, with both the CXCL12-induced Gi-directed activity of CXCR4 (**Fig. 2C**) and constitutive Gi-directed activity of SMO (**Fig. 2D**) observed as the suppression of FSK response. Cell pretreatment with Vismodegib inhibited SMO-mediated suppression of FSK-induced cAMP (**Fig. 2D**), indicating that Vismodegib reduces the constitutive Gi-directed activity of SMO and thus, acts as an inverse agonist. Importantly, cell pretreatment with PTX abrogated both the constitutive activity of SMO and CXCL12-induced activity of CXCR4 (**Fig. 2B-D**), confirming that the observed suppression of FSK-stimulated cAMP downstream of these receptors is indeed mediated by Gi. These temporal trends can be averaged over several consecutive reads and represented as a bar graph reflecting the FSK response (**Fig. 2E**) of respective cell samples; this representation illustrates that FSK-stimulated cAMP is often (but not always) suppressed in SMO-but not in CXCR4-expressing cells, compared to cells with no exogenously transfected receptor. Interestingly, exogenous expression of ACKR3 reproducibly leads to the suppression of FSK-stimulated cAMP and thus creates an impression of constitutive Gi-directed activity similar to SMO. However, the key difference is that SMO-mediated constitutive suppression of FSK-stimulated cAMP is abrogated by cell pretreatment with PTX, whereas ACKR3-mediated suppression remains unchanged. The lack of PTX sensitivity is evidence of ACKR3 not constitutively coupling to Gi, which contradicts findings of [78]. The conversion of the data into the degree of FSK response suppression relative to PTX-treated cells provides a definitive representation clearly indicating constitutive Gi-directed activity of SMO but not any other studied receptor (**Fig. 2F**). Similarly, ligand-induced alteration of the FSK response, compared to the buffer-treated condition, confirms the agonist efficacy of CXCL12 towards CXCR4 and the inverse agonism of Vismodegib towards SMO (**Fig. 2G**). However, no response is observed in ACKR3 transfected cells stimulated with its inverse agonist VUF16840 [96,97], or its agonist CXCL12, consistent with prior literature [74–77] but in disagreement with [79,80].

**Figure 2.**
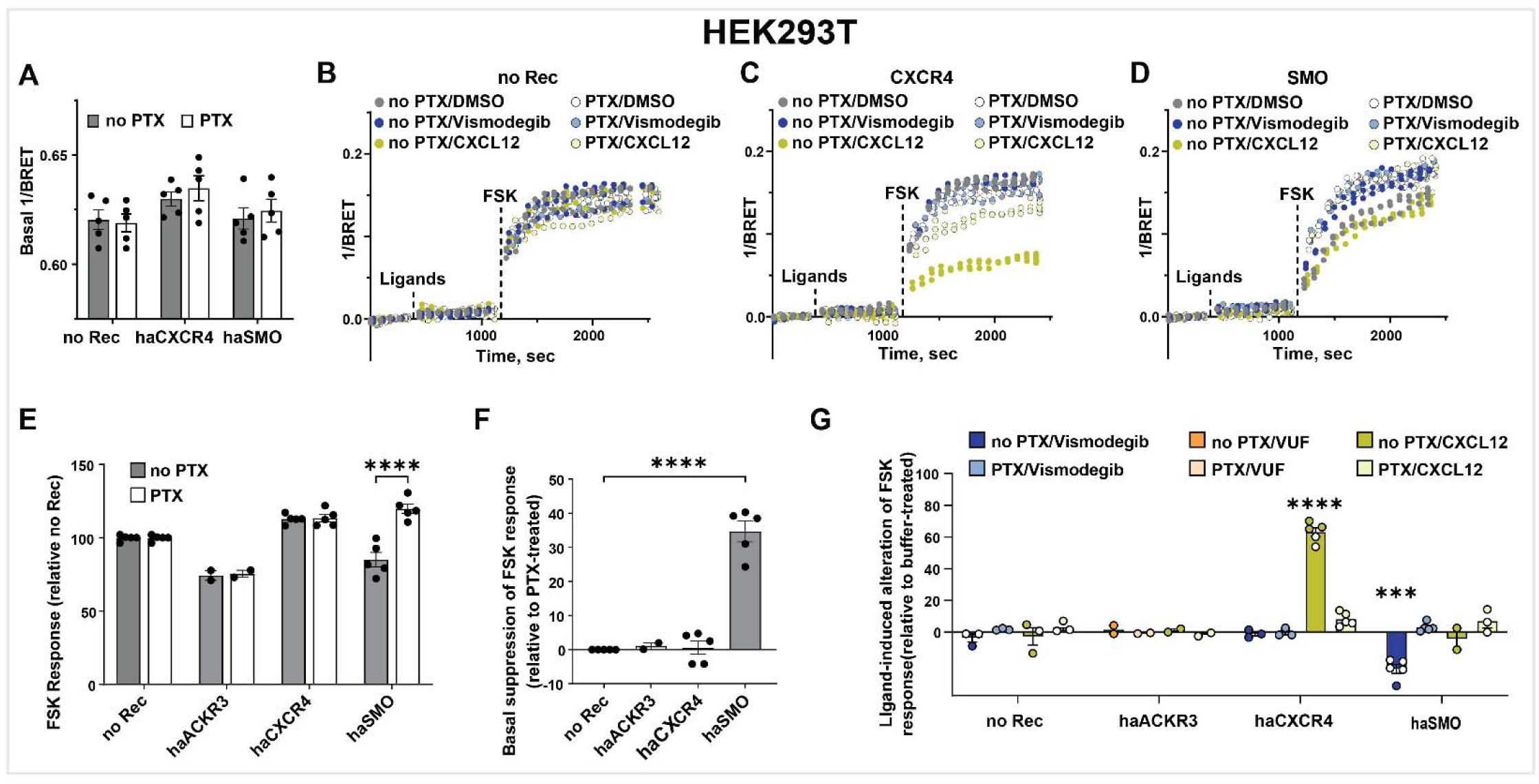
Detection of constitutive and agonist-induced activity of GiPCRs in HEK293T cells using the CAMYEL cAMP biosensor. (**A**) Basal CAMYEL 1/BRET ratios (indicative of intracellular cAMP concentration) in HEK293T cells transfected with no receptor (no Rec), haCXCR4 or haSMO and pretreated or not with PTX. (**B-D**) Time course of changes in 1/BRET ratios in cells upon the addition of agonists or antagonists (‘ligands’) followed by the addition of FSK in HEK293T cells transfected with an empty vector (**B**), haCXCR4 (**C**) or haSMO (**D**). (**E**) Quantification of FSK response of ACKR3-, CXCR4-and SMO-expressing HEK293T cells pretreated or not with PTX. (**F**) Basal suppression of FSK response in ACKR3-, CXCR4- or SMO-expressing HEK293T, quantified relative to the same cells pretreated with PTX. (**G**) Quantification of ligand-induced changes in the suppression of FSK-stimulated cAMP in HEK293T cells expressing ACKR3, CXCR4, or SMO, with and without PTX pretreatment, relative to the same cells treated with buffer. Data in (**A**) and (**E-G**) are from n=3 or more independent experiments (n=2 for ACKR3) performed on different days. Data in (**B-D**) depict technical replicates from one representative experiment. Data in (**A**) and (**E-G**) are plotted as interleaved scatter of individual biological replicates; bar heights reflect the replicate mean and error bars reflect SEM. Significance values in (**E-F**) were determined by two-way ANOVA with Bonferroni’s correction for multiple comparisons. Significance values for **G** were calculated by three-way ANOVA with Bonferroni’s correction for multiple comparisons. ∗P < 0.05, ∗∗P < 0.01, and ∗∗∗∗P < 0.001. FSK, forskolin; PTX, Pertussis Toxin; no Rec, no receptor; sec, seconds.

Collectively, these data confirm the ligand-induced Gi-directed activity of CXCR4, constitutive Gi-directed activity of SMO, and the lack of both types of Gi-directed activity of ACKR3 in HEK293T cells, all through the use of the BRET-based CAMYEL cAMP reporter. It also establishes the degree of FSK response suppression relative to PTX-treated cells as a robust way of assessing constitutive activity of Gi-coupled receptors. Ligand-induced alterations of such activity (such as inverse agonism of Vismodegib at SMO) may also be used to demonstrate constitutive Gi-directed signaling; however, such ligands may be unavailable for orphan receptors [38,59,68]; thus studies in the presence and absence of PTX may be a critical (or only) tool for the discovery of constitutive Gi-directed activity of said receptors.

### Dynamic range of the CAMYEL cAMP assay varies between cell lines and requires cell-line-specific optimization

Encouraged by the successful application of the CAMYEL cAMP assay to studies of constitutive and ligand-induced Gi-directed activity of different receptors in HEK293T cells, we turned our attention to the second cell line, HeLa. Unfortunately, direct transfer of HEK293T CAMYEL BRET conditions to HeLa did not produce a sufficiently high dynamic range or signal-to-noise ratio (**Supp. Fig. 2A**) indicating the need for cell-line specific optimization.

To improve the overall luminescence emission and BRET signal-to-noise ratio in HeLa cells, we replaced the conventional CAMYEL sensor containing circularly permuted (cp) mCitrine and humanized Renilla luciferase (RLuc), with an enhanced brightness biosensor that contains two point mutations in the luciferase component, effectively converting the luciferase into RLuc2 = Rluc(C124A/M185V), similar to [98]. This modification increased luminescence output ∼3-4 fold in both HEK293T and HeLa cells (**Supp. Fig. 2B-C**); therefore, the modified CAMYEL-V2 was used for all subsequent experiments involving HeLa. For experiments with HEK293T cells, the original CAMYEL sensor was used because it provided sufficient luminescence output and signal-to-noise ratio.

Despite the increased signal-to-noise ratio, the absolute magnitude of the FSK response in HeLa cells remained very low, approximately 10% of the response observed under similar conditions in HEK293T cells (**Supp. Fig. 2D**). Therefore we next sought ways to promote a measurable and sustainable FSK response in HeLa cells. First we hypothesized that the low FSK response was due to rapid degradation of cAMP by phosphodiesterases, and tested if it could be improved by cell pre-incubation with isobutylmethylxanthine (IBMX), a non-selective phosphodiesterase inhibitor. Pretreating HeLa cells with IBMX did indeed significantly enhance the overall FSK response (**Fig. 3A**); moreover, the enhanced FSK response improved the signal-to-noise ratio and helped identify specific agonist and inverse agonist responses in CXCR4-and SMO- transfected cells, respectively (**Fig. 3B**). IBMX also improved the FSK response in HEK293T cells (**Fig. 3C**); however, following baseline correction, the magnitude of the CXCL12 agonist and Vismodegib inverse agonist responses actually decreased in IBMX-treated CXCR4- and SMO-transfected HEK293T cells, respectively, compared to no-IBMX (**Fig. 3D**). In other words, by inhibiting the ability of HEK293T cells to degrade cAMP, IBMX appeared to **mask** the constitutive and agonist-induced GiPCR activity in HEK293T cells. This indicates that despite the ability of IBMX to increase the overall FSK response, its utility in the detection of constitutive and ligand-dependent Gi-directed activity of GPCRs varies between different cell lines.

**Figure 3.**
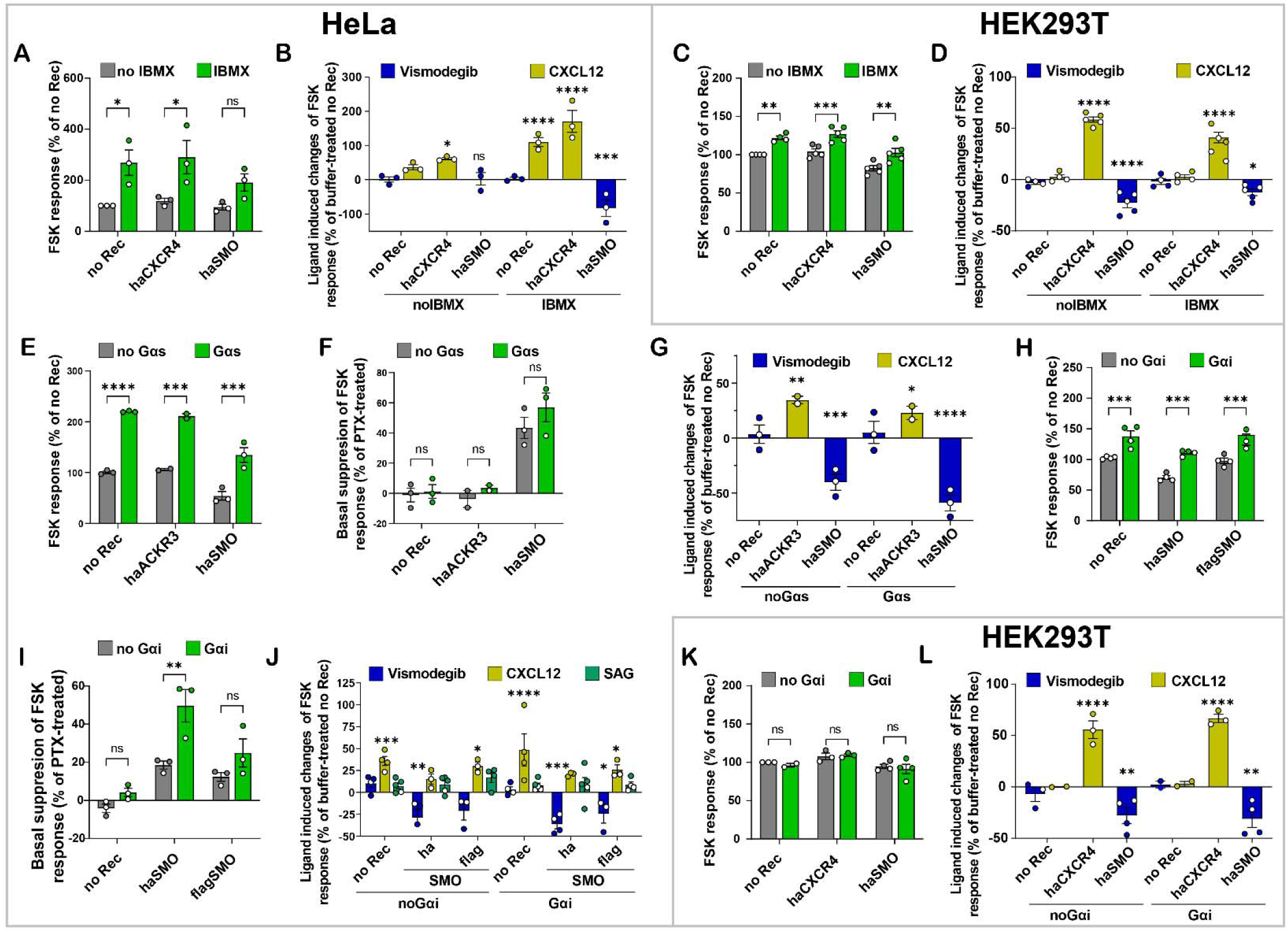
Cell line-specific optimization of the dynamic range of the CAMYEL cAMP assay. (**A**, **C**) FSK response in HeLa (**A**) or HEK293T (**C**) cells expressing the indicated receptors and pretreated or not with IBMX. (**B**, **D**) CXCL12-and Vismodegib-induced changes in FSK response in HeLa (**B**) or HEK293T (**D**) cells exogenously expressing the indicated receptors, relative to buffer-treated no-receptor cells, in the presence or absence of IBMX. (**E**) FSK response in HeLa cells expressing different receptors with and without co-transfected Gαs. (**F**) Basal suppression of FSK response in HeLa cells expressing haACKR3 or haSMO, relative to the same cells pretreated with PTX, with and without co-transfected Gαs. (**G**) Ligand-induced changes in FSK response in HeLa cells transfected with haACKR3 or haSMO with and without co-transfected Gαs, relative to the same cells treated with buffer. (**H**) FSK response in HeLa cells expressing the indicated SMO constructs with and without co-transfected Gαi. (**I**) Basal suppression of FSK response in HeLa cells expressing the indicated receptors, relative to the same cells pretreated with PTX, with and without cotransfected Gαi. (**J**) Ligand-induced changes in FSK response in HeLa cells transfected with the indicated SMO constructs with and without co-transfected Gαi, measured relative to buffer-treated no-receptor cells. (**K**) FSK response in HEK293T cells exogenously expressing haCXCR4 or haSMO with and without co-transfected Gαi. (**L**) Ligand-induced changes in FSK response in HEK293T cells transfected with haCXCR4 or haSMO with and without co-transfected Gαi, measured relative to control buffer-treated no-receptor cells. Data in **A-L** are from 3 or more independent experiments (n=2 for ACKR3) performed on different days. All graphs are plotted as interleaved scatter of individual biological replicates, with bar heights representing replicate means and error bars representing SEM. Significance values were determined by two-way (**A**, **C**, **E-F**, **H-I**, **K**) or three-way (**B**, **D**, **G**, **J**, **L**) ANOVA with Bonferroni’s correction for multiple comparisons. ∗P < 0.05, ∗∗P < 0.01, ∗∗∗P < 0.005 and ∗∗∗∗P < 0.001.

Because FSK sensitivity (as well as basal cAMP levels) have been previously reported to be increased by exogenous expression of Gs-coupled receptors [99,100], we next hypothesized that HeLa FSK response could also be enhanced by cotransfection of Gαs. This indeed proved to be correct: even in the absence of IBMX, HeLa cells transfected with Gαs (along with CAMYEL and receptors of interest) exhibited a change in FSK-stimulated inverse BRET that was approximately two-fold higher than in non-Gαs-transfected cells (**Fig. 3E**). However, the magnitude of FSK response suppression by SMO (compared to PTX-treated cells) was not significantly altered in Gαs-expressing cells (**Fig. 3F**). Similarly, the magnitudes of the agonist (CXCL12) response in haACKR3-transfected HeLa cells and inverse agonist (Vismodegib) in haSMO-transfected HeLa cells were not significantly altered with Gαs-cotransfection, although such cotransfection made these responses less and more significant, respectively (**Fig. 3G**). Noting a robust CXCL12 response in haACKR3- transfected HeLa, we acknowledge that such response could also be attributed to endogenous CXCR4 (see e.g. **Fig. 3B**); we further explore this possibility below. As expected, haACKR3-transfected cells did not demonstrate any constitutive cAMP suppression as evidenced by +/- PTX comparison (**Fig. 3F**). Altogether, this indicates that co-transfection of Gαs does not provide clear advantages for the optimization of detection of Gi-directed receptor activity in the CAMYEL assay in HeLa. The effects of Gαs transfection in HEK293T were not tested.

As a third and complementary strategy for optimizing the detection of Gi-directed constitutive and ligand-induced receptor activity, we tested the cotransfection of Gαi. To take advantage of the prior finding about the role of IBMX, these experiments were done with IBMX in HeLa cells and without IBMX in HEK293T cells. Interestingly and unexpectedly, Gαi co-transfection also improved the FSK sensitivity of HeLa cells (**Fig. 3H**), and concurrently improved the magnitude of the constitutive suppression of the FSK response by SMO (compared to the same cells pretreated with PTX, **Fig. 3I**). Magnitudes of agonist and inverse agonist responses were not significantly altered although some of the responses became more significant (**Fig. 3J**). The effects were consistent between two different variants of SMO constructs N-terminally tagged with HA-or Flag-tags and cloned in two expression vectors (pGEN and pVLAD6 - see **Methods**) (**Fig. 3H-J**), proving receptor specificity. By contrast with HeLa, Gαi co-transfection of HEK293T did not lead to any measurable enhancement of FSK response or the detection of Gi-directed activity of the two tested receptors (**Fig. 3K-L**).

Collectively, these data suggest that the BRET-based CAMYEL cAMP biosensor assay requires cell-line-specific optimization, and that in at least some cell lines (exemplified here by HeLa), the use of IBMX and the co-transfection of Gαi can enhance both the FSK response and the dynamic range of Gi-mediated suppression of such responses. By contrast, the same approaches may have no effect or even be detrimental in other cell lines (exemplified here by HEK293T).

### A step closer: BRET between GRK3ct and G**βγ** identifies constitutive and ligand-dependent activity of GiPCR

Having established the utility and optimal conditions for the characterization of Gi-directed receptor activity through measurements of the second messenger cAMP, we next turned our attention to a more receptor-proximal assay. As described above, agonist-activated GPCRs promote Gα-Gβγ dissociation, and, consequently, the association of Gβγ with its downstream effectors such as GRK2/3. This event can be detected as BRET between GRK3ct-RLuc and mVenus-Gβγ (**Fig. 1D**) [44–48], which we tested in HEK293T and HeLa cells transfected with our panel of receptors (**Fig. 4A-H**).

**Figure 4.**
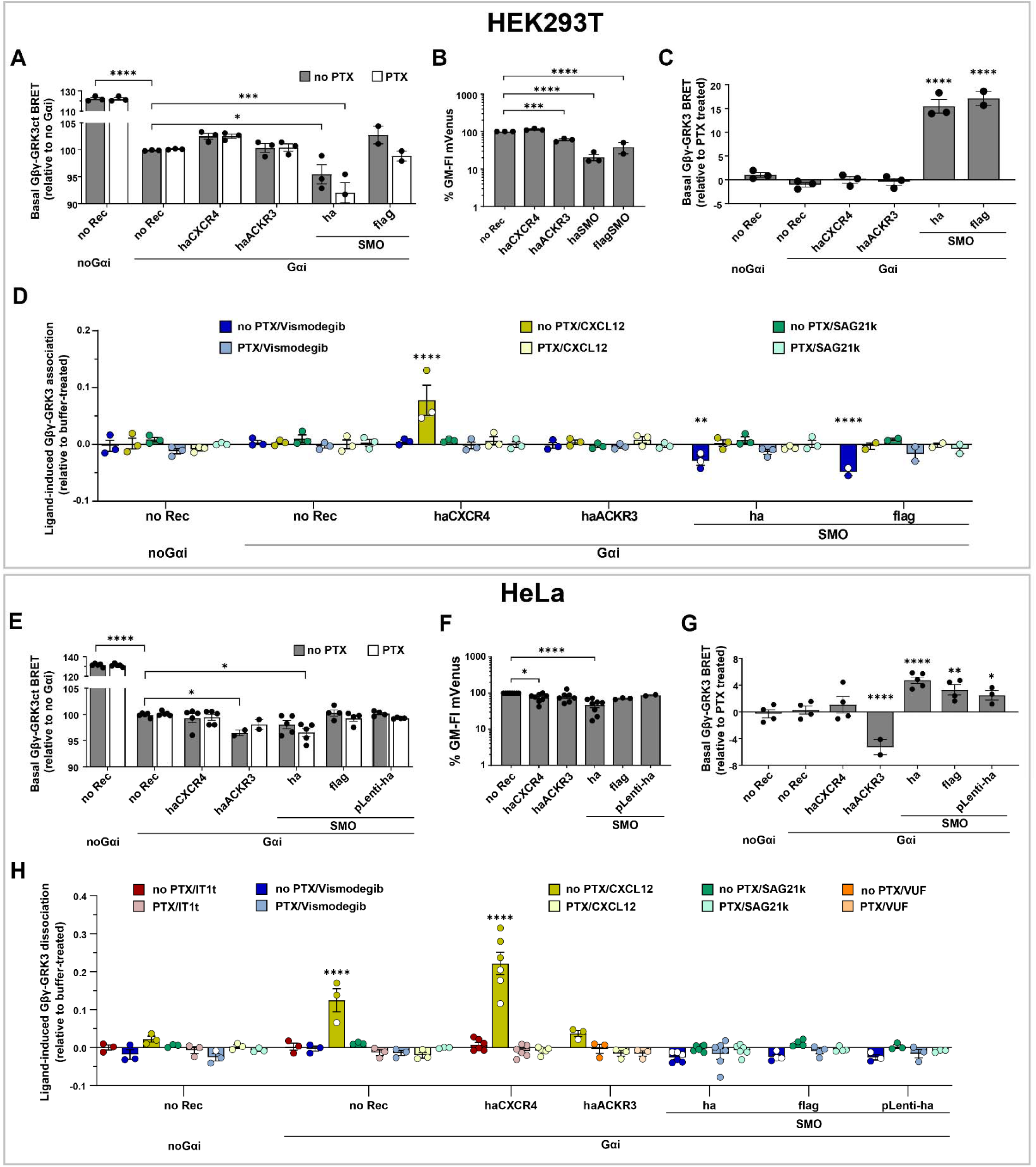
Identification of constitutive and ligand-dependent Gi-directed activity of receptors using BRET between Gβγ and GRK3ct. (**A**, **E**) Basal BRET between Gβγ and GRK3ct in HEK293T (**A**) or HeLa (**E**) cells transfected with the indicated receptors and co-transfected or not with Gαi (BRET ratios are expressed relative to Gαi-transfected no-receptor cells. (**B**, **F**) Expression levels of the acceptor (mVenus-Gβγ) quantified via flow cytometry in HEK293T (**B**) or HeLa (**D**) cells co-expressing the reporter with the indicated receptors. (**C**, **G**) Basal BRET between Gβγ and GRK3ct in HEK293T (**C**) or HeLa (**G**) cells transfected with the indicated receptors, relative to the same cells pretreated with PTX. (**D, H**) Ligand-induced changes Gβγ-GRK3ct BRET in HEK293T (**D**) and HeLa (**H**) cells expressing the indicated receptors, relative to the same cells treated with buffer. Data in **A-H** are from 2-8 independent experiments performed on different days. All graphs are plotted as interleaved scatter of individual biological replicates, with bar heights representing replicate means and error bars representing SEM. Significance values in **A** and **E** were determined by two-way ANOVA, in **B-C** and **F-G** by one-way ANOVA and in **D**, **H** by three-way ANOVA, and with Bonferroni’s correction for multiple comparisons. ∗P < 0.05, ∗∗P < 0.01, ∗∗∗P < 0.005 and ∗∗∗∗P < 0.001.

We first noticed that cell co-transfection with GRK3ct-RLuc and mVenus-Gβγ results in a very high level of constitutive association between them, both in HEK293T (**Fig. 4A**) and in HeLa (**Fig. 4E**), indicative of the inability of endogenous Gαi to sequester the excess transfected Gβγ and keep it in the inactive heterotrimer form. Consistent with this explanation, co-transfecting additional exogenous WT Gαi significantly suppressed GRK3ct-RLuc/mVenus-Gβγ basal association (**Fig. 4A,E**). Unexpectedly, the GRK3ct-RLuc/mVenus-Gβγ BRET was even more suppressed in HEK293T and HeLa transfected with haSMO, as well as in HeLa transfected with haACKR3 (**Fig. 4A,E**). This direction of change in BRET was unexpected because taken at a face value, it suggests that haSMO promotes the formation of the inactive Gαi/mVenus-Gβγ heterotrimers, contrary to its ability to constitutively activate Gαi cAMP signaling (**Fig. 3F, I**). After careful examination, we attributed this effect to variations in reporter expression, as haSMO strongly suppressed the expression of mVenus-Gβ1 as assessed by flow cytometry, both in HEK293T (**Fig. 4B, Supp. Fig. 3A**) and in HeLa (**Fig. 4F, Supp. Fig. 3B**). This illustrates that the GRK3ct-RLuc/mVenus-Gβγ BRET assay may generate artifacts that require caution in the interpretation and can be dissected with the help of flow-cytometry-based assessment of acceptor expression levels.

Importantly, measuring the GRK3ct-RLuc/mVenus-Gβγ BRET in cells co-transfected with Gαi and receptors of interest with and without pretreatment with PTX revealed the expected patterns: in SMO-transfected cells, but not in cells transfected with CXCR4, ACKR3, or no exogenous receptors, PTX pretreatment reduced the GRK3ct-RLuc/mVenus-Gβγ association indicating the reduction in free Gβγ and restoration of inactive Gαi/Gβγ heterotrimers. The comparative +/- PTX representation of this data (**Fig. 4C,G**) clearly illustrates the constitutive Gi-directed activity of SMO in several N-terminally tagged versions and plasmid backbones, and in both HEK293T and HeLa cells. It also establishes the lack of such activity for all other tested receptors including ACKR3.

Despite the above caveats, the GRK3ct-RLuc/mVenus-Gβγ BRET association assay proved suitable for the detection of ligand-induced activation or suppression of Gi-directed activity of the tested receptors. In particular, CXCL12 stimulation led to a robust increase in GRK3ct/Gβγ association in CXCR4-transfected but not any other cells, whereas pretreatment with Vismodegib decreased such association in SMO-transfected HEK239T (**Fig. 4D,H**). Pretreatment with PTX eliminated these responses (**Fig. 4D,H**), confirming that the responses are Gi-mediated. Stimulation of SMO-expressing cells with the small-molecule SMO agonist SAG21k [101] produced no change in GRK3ct/Gβγ association (**Fig. 4D,H**), likely because the high levels of SMO constitutive activity in this setup make it unresponsive to further activation by an agonist. The trends held across multiple tested variants of N-terminally HA- or Flag-tagged SMO in pGEN, pVLAD6 and pLenti vectors (see **Methods**) in HEK293T (**Fig. 4D**) but did not reach statistical significance in HeLa.

Taken together, this data establishes the utility of the GRK3ct-RLuc/mVenus-Gbg BRET assay for the detection of constitutive and ligand-induced Gi-directed activity of GPCRs. Being more “proximal” to receptor activation than the CAMYEL cAMP measurement, this assay could be transferred from HEK293T cells to HeLa cells almost without optimization. However, our results also revealed two important caveats (the need for co-transfecting Gαi and BRET signal alteration through the suppression of the reporter expression) that could lead to possible artifacts and mis-interpretation of the results. We also outlined a system of comparisons and controls (most importantly, measurements +/- PTX) that are needed to overcome these caveats.

### As close as it gets: BRET between G**α**i and G**βγ** is most sensitive towards constitutive and ligand-dependent activity of GiPCR

Next we sought to evaluate the third BRET-based assay outlined in **Fig. 1C**: that measuring the dissociation between a donor (RLuc2)-tagged Gαi and acceptor (mVenus)-tagged Gβγ. Because among the studied assays, this assay is most proximal to the receptor activation event in live cells (**Fig. 1A**), we also expected it to be the most sensitive and applicable across both cell lines tested.

As with the GRK3ct-RLuc/mVenus-Gbg association assay described above, this assay requires co-transfection of multiple plasmids. The resulting BRET is affected by the ratio of expression of the BRET donor (here Gαi(91)-RLuc2) and the BRET acceptor (here mVenus-Gβγ); to reduce experiment-to-experiment variation, it is recommended that the acceptor is expressed in large excess of the donor. This is usually achieved by adjusting the amounts of transfected DNA; the amount of the donor is selected as low as possible to allow for a large excess of the acceptor at constant total transfected DNA, while also producing sufficiently bright luminescence and an interpretable BRET ratio in the experiment. As an alternative to this approach, the reporter has also been developed in the form of a single tricistronic vector, pIRES Gβ-2A-cpVenus-Gγ2-Gαi- Nluc [41,43]; this enables transient simultaneous delivery of all three reporter genes in controllable ratios.

To measure the constitutive and ligand-induced activity of the receptors of interest, we used both the three-plasmid and the single tricistronic [41] versions of the Gαi/Gβγ BRET reporter in HEK293T cells, with very consistent results between the two, but only the three-plasmid version in HeLa cells. This is because despite multiple attempts and contrary to the original publication [41], we were not able to establish the tricistronic vector assay in HeLa cells due to an extremely rapid drop of luminescence that prevented accurate BRET measurements over a sufficient period of time (*data not shown*).

As with the GRK3ct-RLuc/mVenus-Gβγ assay above, the basal BRET between donor-tagged Gαi and acceptor-tagged Gβγ was strongly dependent on the transfected receptor in HEK293T (**Fig. 5A**). We also confirmed that not only the constitutive activity but also the suppression of acceptor expression by selected receptors (**Supp. Fig. 3A**) contributes to the observed variations in BRET. Therefore, in this assay too, caution should be exercised when interpreting the basal BRET signal. This was the case even when the tricistronic reporter [41] was used in HEK293T cells (**Fig. 5A, Supp. Fig. 3A**), even though this version of the reporter was designed to alleviate artifacts associated with reporter component expression differences [41].

**Figure 5.**
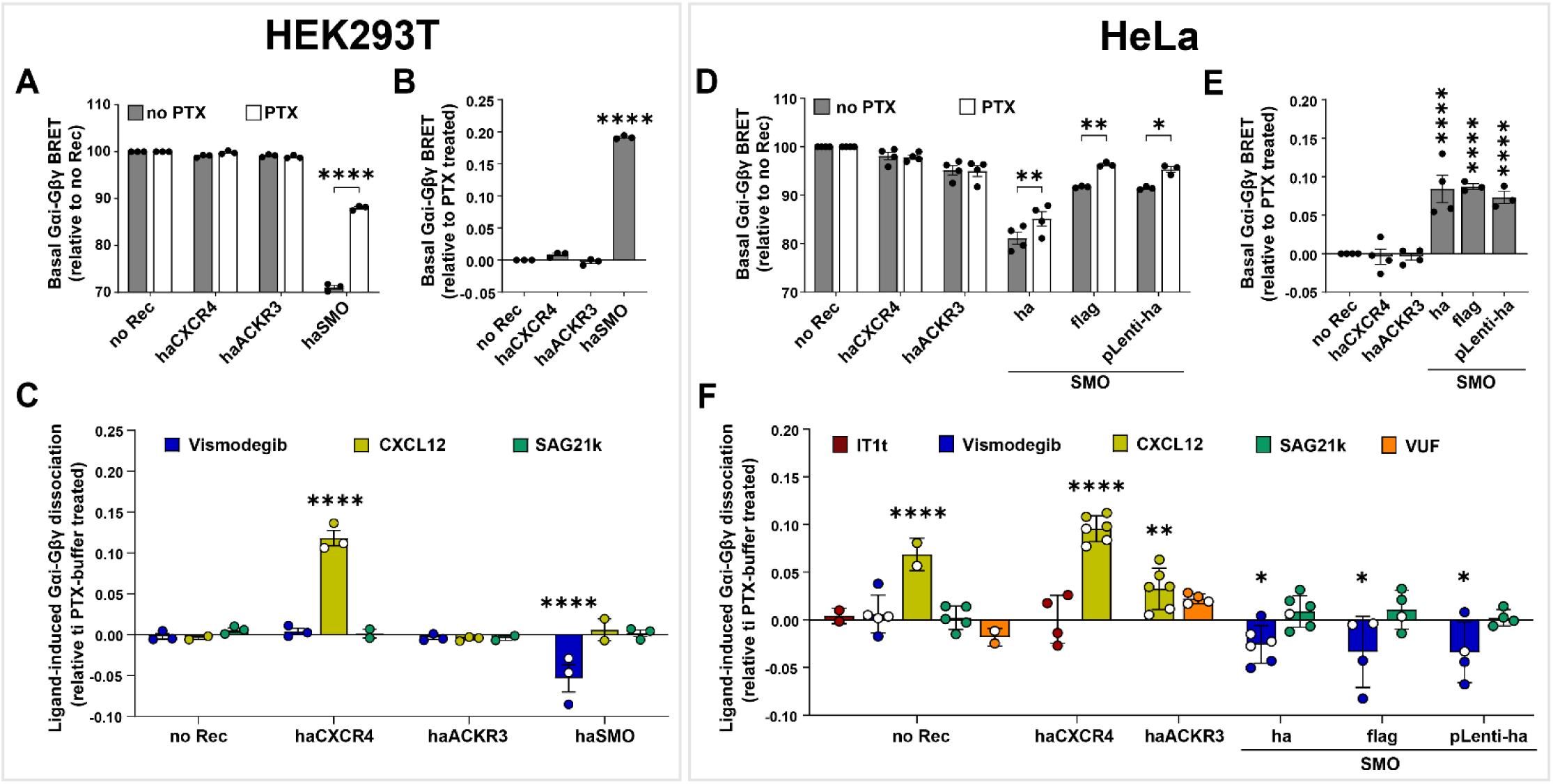
Identification of constitutive and ligand-dependent activity of receptors using BRET between Gαi and Gβγ subunits. (**A, D**) Basal BRET between Gαi and Gβγ in HEK293T (**A**) or HeLa (**D**) cells transfected with the indicated receptors and pretreated or not with PTX. (**B, E**) Basal Gαi/Gβγ dissociation in HEK293T (**B**) or HeLa (**E**) cells transfected with the indicated receptors, expressed relative to the same cell pretreated with PTX. (**C, F**) Ligand-induced changes in Gαi-Gβγ dissociation in HEK293T (**C**) or HeLa (**F**) cells transfected with the indicated receptors, relative to the same cells pretreated with PTX and treated with buffer. Data are from 3 or more independent experiments performed on different days. Graphs are plotted as interleaved scatter of individual biological replicates; bar heights represent replicate means and error bars represent SEM. Significance values were determined by two-way ANOVA (**A-B, D-E**) or three-way ANOVA (**C**, **F**) with Bonferroni’s correction for multiple comparisons. ∗P < 0.05, ∗∗P < 0.01 and ∗∗∗∗P < 0.001.

Despite this caveat, we were again able to robustly separate reporter expression-associated BRET alterations from real manifestation of receptor constitutive activity with the use of PTX. Cell pretreatment with PTX partially eliminated the BRET suppression in SMO-transfected cells but not in cells transfected with other receptors (including ACKR3) or no exogenous receptor (**Fig. 5A-B**).

CXCL12 stimulation led to a significant increase in Gαi/Gβγ dissociation in CXCR4-expressing HEK293T cells; while pretreatment with Vismodegib led to a reduction in Gαi/Gβγ dissociation in SMO-expressing cells, due to elimination of the receptor’s constitutive activity (i.e. Vismodegib again acted as an inverse agonist). Both of these responses were abolished in cells pretreated with PTX. No response was observed when SMO-expressing cells were stimulated with SMO agonist SAG21k (**Fig. 5C**), presumably due to the high constitutive activity of SMO that makes it unresponsive to further activation with agonists.

To confirm that the observations with this reporter are consistent regardless of cell type, the assay was repeated in HeLa cells. As with HEK293T cells, basal Gαi-RLuc/mVenus-Gβγ BRET in HeLa was strongly suppressed by co-transfection of HA-tagged SMO in the pGEN vector, due to the suppression of reporter expression by this receptor construct (**Fig. 5D, Supp. Fig. 3A**). With all three SMO constructs (HA-tagged SMO in pGEN, Flag-tagged SMO in pVLAD6, or HA-SMO in pLenti), but not with any other receptors including ACKR3, the suppression was blocked by cell pretreatment with PTX (**Fig. 5E**): this indicates that PTX works on luciferase-tagged Gαi similar to its effects on endogenous or exogenously transfected WT Gαi, and that the observed SMO-mediated suppression of BRET is construct-and tag-independent but Gi-mediated.

As with HEK293T cells, Gαi-Gβγ dissociation was significantly enhanced in haCXCR4-expressing HeLa cells stimulated with CXCL12, and inhibited in SMO-expressing HeLa cells (HA-, Flag-or pLenti-HA-versions) treated with Vismodegib (**Fig. 5F**). No evidence of constitutive Gαi-Gβγ dissociation, or of modulation of such dissociation by SMO ligands (Vismodegib or SAG21k), was observed in control cells without exogenously transfected receptors. Interestingly, here again, CXCL12 stimulation produced a small increase in Gαi-Gβγ dissociation in ACKR3-expressing HeLa cells (**Fig. 5F**), even though they were pretreated with IT1t to eliminate endogenous CXCR4 signaling. However, because the CXCL12 response was also observed in control HeLa with no transfected receptor (**Fig. 5F**), we concluded that it was still attributable to endogenous CXCR4 whose signaling was not fully abrogated by IT1t. ACKR3 antagonist VUF-16840 had no effect in either control or ACKR3-expressing cells when used alone (**Fig. 5F**). Pretreating cells with VUF-16840 did not alter the response to subsequently applied CXCL12 (**Supp. Fig. 5**) suggesting that the response was not mediated by ACKR3.

These findings confirm that BRET-based measurements of Gαi/Gβγ dissociation provide a robust method for detecting constitutive or ligand-dependent activity of various receptors towards Gi. As expected from the proximity of the reporter to the receptor activation event, the signals were clearer and more consistent, e.g. between all three constructs of SMO in HeLa (**Fig. 5F**). As with the other assays described above, the +/- PTX comparison provided a critically important tool for separating real Gi-directed constitutive activity from artifacts associated with variations in reporter expression, which could be especially important in characterizing orphan receptors with no known direct or inverse agonists.

### BRET between G**α**i and G**βγ** reveals temporal profiles of Gi activation by SMO and its regulation by PTCH1

We next employed the Gαi/Gβγ dissociation assay to characterize the ability of PTCH1 to modulate the Gi-directed SMO activity. PTCH1 is a 12TM protein with an intrinsic sterol flippase activity [102,103] and a central component of the Hedgehog signaling pathway [104] where PTCH1 suppresses constitutive SMO signaling. The function of PTCH1 as a regulator of SMO activity was initially described in the context of canonical primary-cilia-restricted Hedgehog signaling towards the Gli1/2/3 transcription factors [105]; it was also later shown to regulate constitutive SMO signaling towards Gi in HEK293 cells [60,62,69,105].

To explore the modulation of constitutive activity of SMO by PTCH1, Gαi/Gβγ dissociation was measured by BRET in HEK293T cells transiently transfected with SMO (HA-or Flag-tagged SMO in pGEN, pVLAD6, or pLenti) and co-transfected or not with PTCH1b (an engineered variant of mouse PTCH1 with improved expression compared to WT PTCH1 [62,106]).

Gi-directed constitutive activity of SMO, measured as the difference in Gαi-RLuc2/mVenus-Gβγ BRET between PTX-treated and non-PTX-treated HEK293T cells, was significantly decreased in the presence of PTCH1 compared to the no-PTCH1 condition for all three SMO constructs tested (**Fig. 6A**). No evidence of constitutive suppression of Gαi-Gβγ association, or of modulation by PTCH1, was detected in control cells without exogenous receptors (**Fig. 6A**).

**Figure 6.**
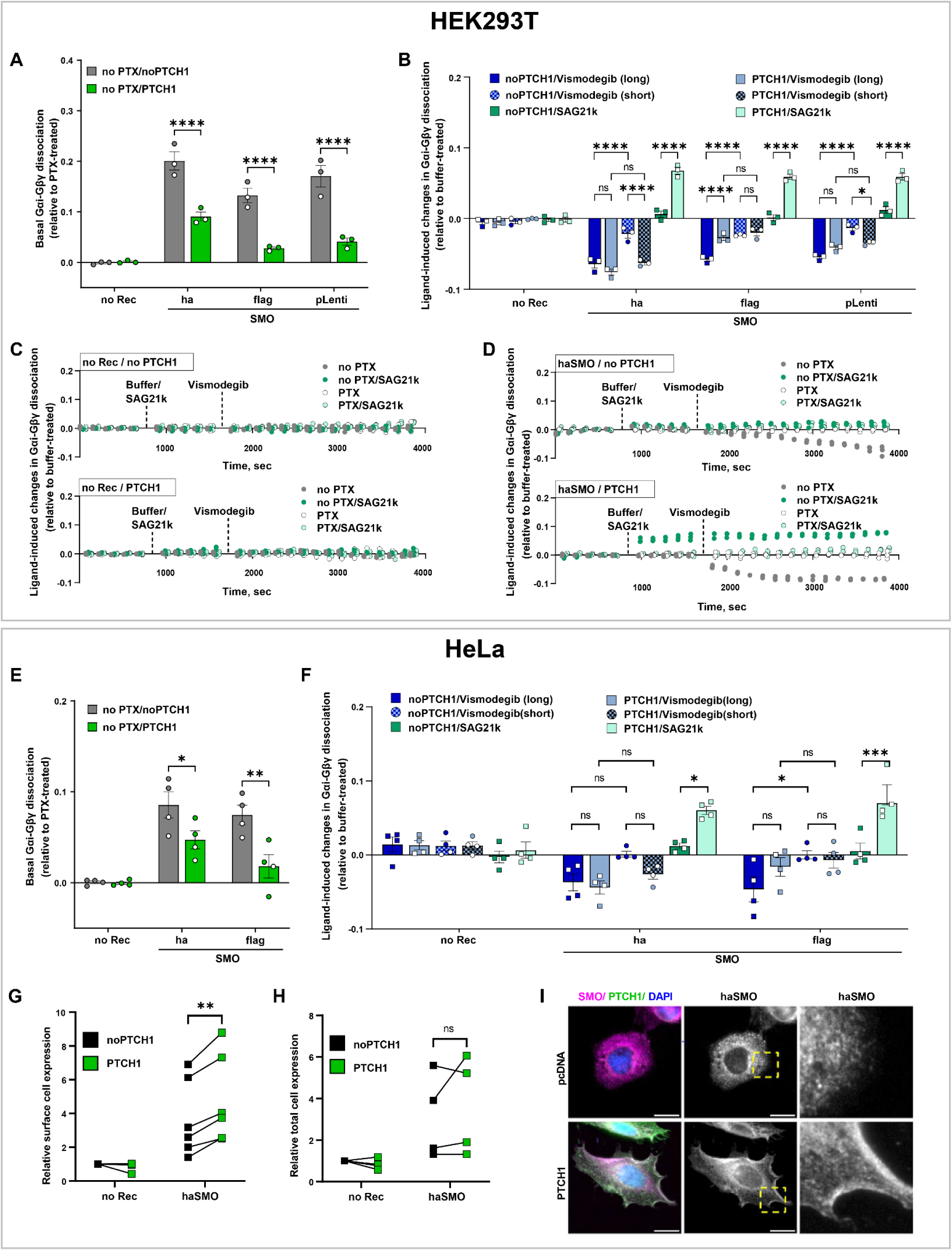
PTCH1 increases SMO sensitivity to ligands by increasing its surface expression. (**A, E**) Basal Gαi-Gβγ dissociation in HEK293T (**A**) and HeLa (**E**) transfected with the indicated SMO constructs and cotransfected or not with PTCH1, relative to the same cells pretreated with PTX. (**B, F**) Ligand-dependent changes in Gαi-Gβγ dissociation in HEK293T (**B**) or HeLa (**F**) cells transfected with the indicated SMO constructs and co-transfected or not with PTCH1. Gαi-Gβγ dissociation is expressed relative to the same cells pretreated with buffer. “Short” ligand treatment corresponds to 10-15 min and “long” to 30-40 min of ligand exposure. (**C, D**) Time courses of changes in Gαi-Gβγ dissociation in HEK293T cells transfected (**D**) or not (**C**) with haSMO, cotransfected or not with PTCH1, and pretreated or not with PTX, following the addition of buffer, SAG21k (agonist), or Vismodegib (antagonist). (**G, H**) Surface (**G**) or total (**H**) expression of haSMO, measured by flow cytometry in HeLa cells co-transfected or not with PTCH1. **(II)** Immunofluorescence of cells transfected with haSMO (magenta) and co-transfected or not with PTCH1 (green); nuclei are shown in blue. Scale bars = 20 µm. Data in **A-B** and **E-H** are from 3 or more independent experiments performed on different days. Data in **A-B**, **E- F** are plotted as interleaved scatter of individual biological replicates; bar heights represent replicate means and error bars represent SEM. Data in **C-D** are technical replicates from one representative experiment shown in **B**. Significance values were determined by two-way ANOVA (**A**, **E**) or three-way ANOVA (**B**, **F**) with Bonferroni’s correction for multiple comparisons. Data shown in **G** and **H** were plotted repeated measures, matched values spread across a row. Significance values for those were calculated by multiple paired t-test. ∗P < 0.05, ∗∗P < 0.01, ∗∗∗P < 0.005 and ∗∗∗∗P < 0.001.

Next we studied the impact of PTCH1 co-expression on SMO responses to its ligands. In the presence of PTCH1, all three SMO constructs robustly responded to SMO agonist SAG21k (**Fig. 6B**): this is consistent with prior reports where PTCH1-mediated suppression of SMO constitutive activity makes SMO responsive to agonists [107–109]. Unexpectedly, we also observed that PTCH1 co-transfection enhanced SMO response to short (∼10-15 min) but not longer (∼30-40 min) pretreatment with inverse agonist Vismodegib (**Fig. 6B**). Intrigued by this observation, and capitalizing on the time-resolved nature of the Gαi-RLuc2/mVenus-Gβγ BRET assay, we recorded the time-course of Gαi/Gβγ dissociation in SMO-expressing cells co-transfected or not with PTCH1, pretreated or not with PTX, and stimulated or not with agonist SAG21k or inverse agonist Vismodegib (**Fig. 6C-D**). Non-SMO-expressing cells showed no responses to ligands regardless of PTCH1 co-expression (**Fig. 6C**). SMO-expressing cells contransfected with PTCH1 responded to SAG21k almost immediately (**Fig. 6D**), in agreement with **Fig. 6B** and prior studies [107,108]. Furthermore, consistent with our earlier end-point measurements, both PTCH1-expressing and non-PTCH1-expressing SMO-transfected cells responded to Vismodegib; however, intriguingly, the response was delayed in the absence of PTCH1 and more rapid in PTCH1-transfected cells (**Fig. 6D**). This suggests that PTCH1 sensitizes SMO not only to agonists but also to inverse agonists: an aspect of PTCH1 regulatory function that, to our knowledge, has not been reported previously.

Using the same approach as for HEK293T, the constitutive activity of SMO and its modulation by PTCH1 was examined in the second cell line, HeLa, with consistent results: basal Gα-Gβγ dissociation induced by SMO (HA-and Flag-tagged versions) was significantly reduced when the cells were co-transfected with PTCH1 (**Fig. 6E**). PTCH1-mediated acceleration of responses to inverse agonist Vismodegib was less pronounced in HeLa compared to HEK293T (**Fig. 6F**). As with HEK293T cells, there was a significantly enhanced response to SAG21k in HeLa expressing both SMO and PTCH1 compared to HeLa expressing SMO alone (**Fig. 6F**).

We hypothesized that the acceleration of SMO responses to both agonists and inverse agonists could be mediated by altered subcellular distribution and higher availability of functional SMO at the cell surface in the presence of PTCH1. Indeed, flow cytometry demonstrated a reproducible increase in cell surface levels of haSMO in the presence of PTCH1, compared to non-PTCH1-expressing cells (**Fig. 6G, Supp Fig. 4A**), while total levels did not change significantly (**Fig. 6H, Supp Fig. 4B**). Accordingly, immunofluorescence analysis showed enrichment of haSMO on the membrane of HeLa co-transfected with PTCH1 (**Fig. 6I**). Collectively, these data show that in addition to its well-documented function of suppressing the constitutive activity of SMO, PTCH1 may also enhance SMO responsiveness to inverse agonists by altering its cellular distribution and increasing its levels on the cell surface. The data also illustrates the applicability of the BRET-based Gαi/Gβγ dissociation biosensors for hypothesis generation and characterizing subtle aspects of GiPCR function in a time-resolved manner in various cell types.

## Discussion

In this study, we investigated the applicability of three BRET-based G protein activation reporters for the characterization of constitutive and ligand-dependent Gi-directed activity of GPCRs in live cells. All three assays are directly related to G protein signaling (as opposed to e.g. β-arrestin recruitment, receptor trafficking, or integrated downstream responses such as cell migration or dynamic mass redistribution, DMR [110–112]), but vary in the degree of their “proximity” to the receptor-G protein coupling event.

Detection of constitutive Gi-directed activity of GPCRs is an important and challenging task, especially for orphan receptors without known ligands, and for endogenous contexts and primary cells where the signaling profiles of such receptors can be dramatically different. An infamous example illustrating these challenges is the orphan GPR37L1 that signals to its proposed ligand prosaptide TX14(A) in endogenously expressing cells - primary astrocytes [113,114], - but not in heterologously transfected cells lacking endogenous GPR37L1 such as HEK293T/17, CHO, and NIH-3T3 [68,115]. Because the methods presented in this paper do not rely on receptor tagging, they can be applied to orphan Gi-coupled receptors in systems where (i) robust reporter expression can be achieved and (ii) controls with and without the receptor are available. Here, we used transiently transfected HEK293T and HeLa cells; however, for difficult-to-transfect systems, reporter delivery with the use of lentiviruses and adeno-associated viruses, and the establishment of stably expressing reporter systems, may be necessary. For endogenously expressed receptors, the use of anti-receptor shRNA or CRISPR KO may be necessary to confirm receptor specificity [113]. Experiments with direct effector recruitment to tagged receptors are another way to ensure specificity [29–37]; not only this is achievable in heterologous expression systems but studies have also been published where tags are introduced into endogenous receptors via CRISPR [90]. However, tagging can alter the signaling and trafficking properties of the receptor [28,116,117], and therefore we purposely excluded such assays from the present study.

The most “remote” of the studied assays, CAMYEL [54], detects GiPCR activity through the decrease in the concentration of intracellular cAMP resulting from Gαi-mediated inhibition of AC. The need to measure a decrease (rather than an increase) of cAMP presents a number of challenges and, as we demonstrate, requires optimization for the cell lines of choice. The Gβγ/GRK3ct BRET assay is more proximal but has limited utility at endogenous Gα levels and can generate artifacts via receptor-associated variations in reporter expression levels. Constitutive activation of Gi by GPCRs is best detected through Gαi-Gβγ dissociation assay because it is the most proximal to the receptor and thus provides the highest sensitivity. Accordingly, variations of this biosensor are widely used for GPCR activity characterization in different contexts. For example, Schihada et al. demonstrated the use of their G-CASE (Gα-NLuc/cpVenus-Gγ) BRET biosensors not only in HEK293 cells but also in more physiologically relevant mouse lung epithelial cells (MLE-12) without overexpression of exogenous GPCRs [43]. Similarly, using their pIRES Gβ-2A-cpV-Gγ2-Gα-Nluc biosensor, Matti et al. demonstrated signaling incompetence of the atypical chemokine receptor ACKR4 towards five types of G proteins [41]. Finally, the conceptually similar TRUPATH BRET2 Gαβγ biosensors with optimized positioning of donor and acceptor on the Gα and Gγ subunits have shown excellent dynamic range and enabled the determination of G protein transducerome (receptor activity towards 16 G protein subtypes) for multiple receptors [42].

When it comes to the detection of constitutive activity and inverse agonists are not available, the above assays require properly designed comparisons. In our work, we used comparisons with empty-vector-transfected cells (which is not ideal due to the need to compare BRET measurements between different transfectants) and with the same transfectant pretreated with Gi inhibitor PTX [93–95]. We demonstrated that PTX pretreatment provides a robust way to distinguish constitutively active GiPCRs from receptors that do not possess such activity. However, this approach is only applicable to GiPCRs, and specifically only to those with the canonical constitutive coupling geometry: it may not work for the hypothetical “non-canonical” GPCR/Gi coupling geometries [118]. For Gq-directed activity, in similar contexts, Gαq-specific inhibitors such as UBO-QIC (also known as FR900359 [119]) have been used [52]. We are not aware of similarly acting pharmacological inhibitors for Gαs and Gα12/13.

In the absence of a pharmacological inhibitor for the Gα protein of interest, the design of comparisons for the detection of constitutive GPCR activity can be challenging. To this end, Schihada et al. proposed assessing the relationship between G-CASE BRET and NLuc luminescence in the same cells, and demonstrated the utility of this approach for four GPCRs with intrinsic coupling to different G proteins [43]. Alternatively, Kuravi et al. developed a unique approach, capitalizing on rapamycin-induced proximity between an FKBP-tagged membrane-targeted ‘third-party’ BRET donor and FRB-tagged Gβγ which leads to an increase in BRET between the donor and acceptor-tagged Gα; in this scenario, the comparison in the presence vs absence of rapamycin presents the desired path towards the detection of constitutive GPCR activity [120].

The invention of the enhanced bystander BRET (ebBRET) technology [49,121] greatly diversified the range of molecular events that can be detected without receptor or even G protein tagging [49,51,52]. This could be achieved via bimolecular BRET sensors where the acceptor is targeted to the PM (e.g. rGFP-CAAX) or another subcellular compartment of interest (e.g. rGFP-FYVE for early endosomes), and the donor is a known effector of Gβγ-free, GTP-bound Gα (e.g. Rap1GAP for Gαi/o-GTP, p63-RhoGEF for Gαq/11-GTP, or PDZ-RhoGEF for Gα12/13-GTP and Gs) [49,51,52]. Alternatively, the ‘BERKY’ family of unimolecular ebBRET-based biosensors [50,53] consist of CAAX-NLuc-linker-YFP attached to a known Gαi-GTP peptide binder KB-1753, or to other sensors for different Ga isoforms; these unimolecular biosensors allow for sensitive detection of activation of *endogenou*s G proteins not only in cell lines but also in primary cells such as lentivirally transduced mouse cortical neurons [50,53]. The use of these sensors for the detection of GPCR constitutive activity has not been reported; however, in principle it should be feasible using the same controls as described above.

With the use of the Gαi-Gβγ dissociation assay, we demonstrated the previously documented [60,70,71] ability of PTCH1 to inhibit the constitutive activity of SMO [60,62,69,108,109]. Moreover, PTCH1 promotes SMO localization at the cell surface, thus, making it more responsive not only to agonists but also to antagonists and inverse agonists. While the canonical Hedgehog pathway signaling is from SMO to the GLI transcription factors, the ability of SMO to regulate Gi, and the cAMP-PKA axis is getting increasingly appreciated [62,122–124]. In this context, our findings may shed new light on the mechanisms that regulate SMO-mediated activation of Gi.

Altogether, we demonstrated the use of three BRET-based G protein activation reporters for the characterization of constitutive and ligand-dependent Gi-directed activity of GPCRs in live cells at different levels of “proximity” to the receptor. All three assays are directly related to Gi signaling and can be used in different cell types; however, at least one of them (CAMYEL) required cell-line-specific optimization. By contrast, the Gαi/Gβγ dissociation assay demonstrated the most robust results and utility for the discovery of GPCR signaling profiles.

## Methods

### Reagents and supplies

Commercially available reagents and supplies used in this work are listed in **Table 1**. The plasmids used in the paper are listed in **Table 2**.

**Table 1.**
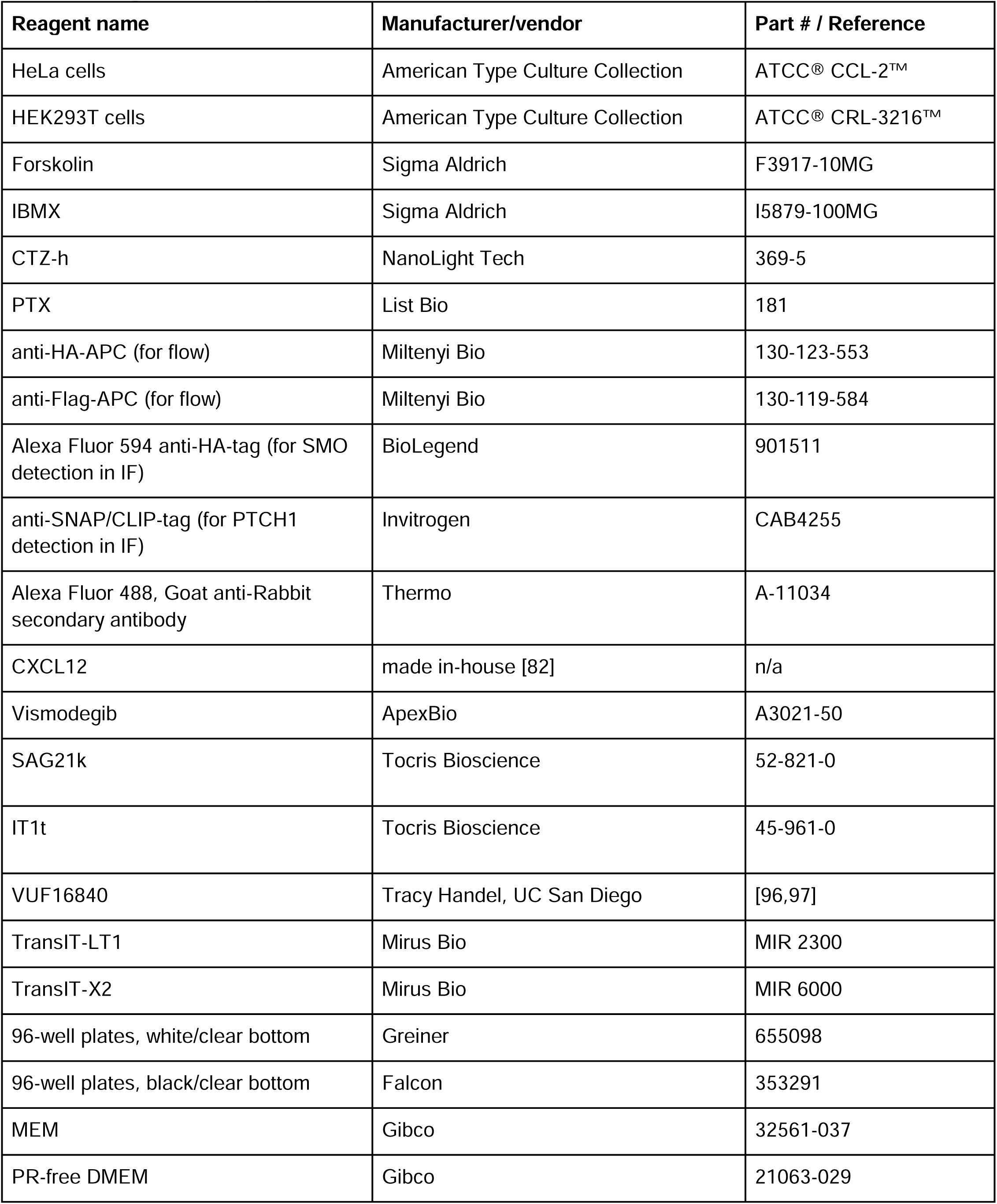
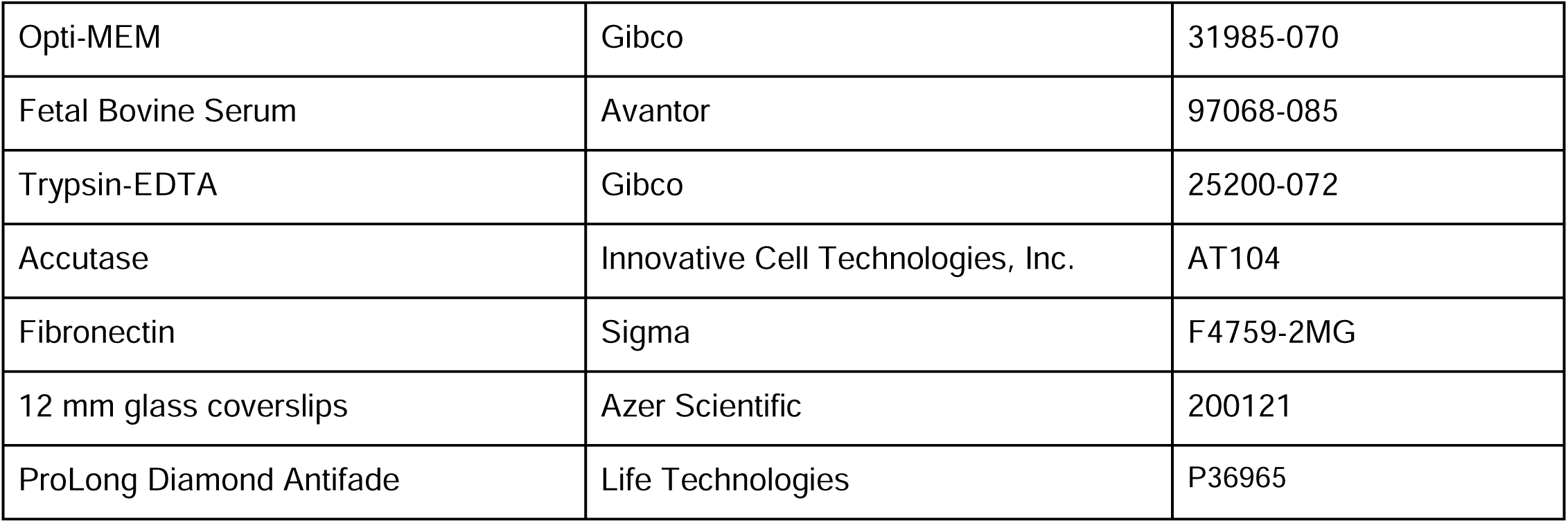
Reagents and supplies.

**Table 2.**
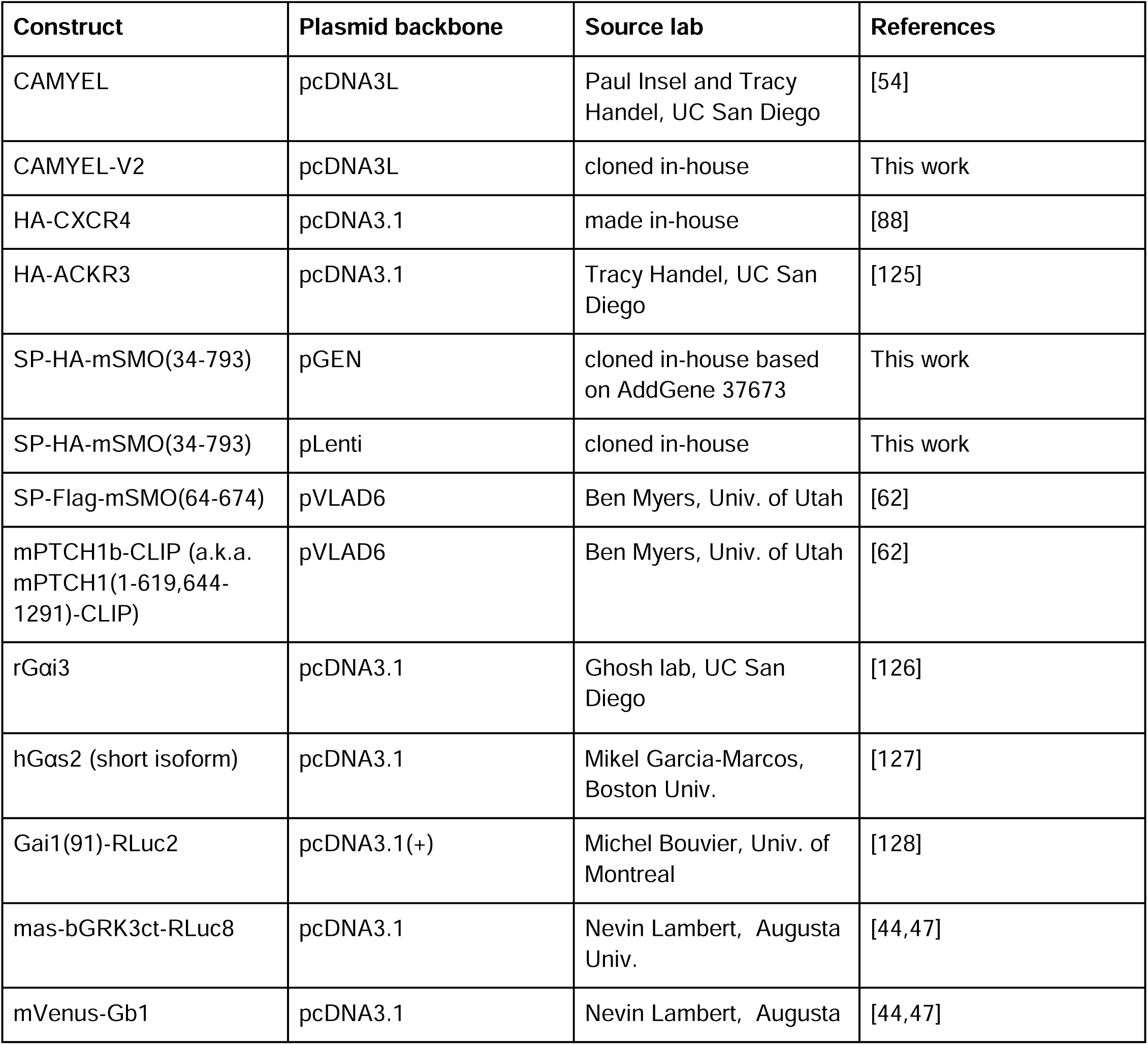
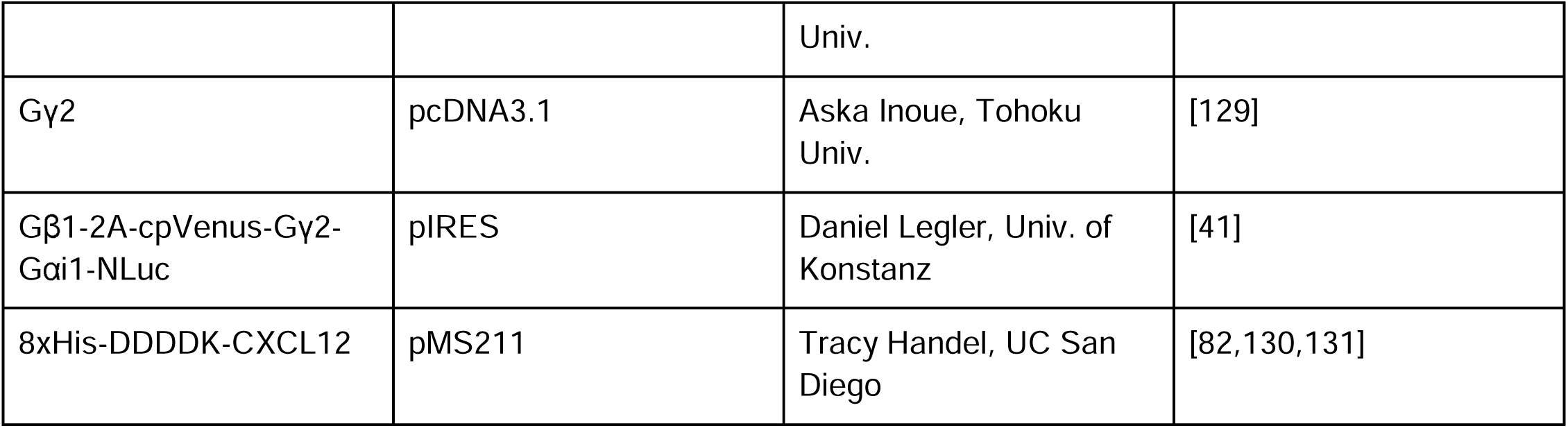
Constructs and plasmids used in this work.

### Cell culture and transfection

HEK293T were cultured in Dulbecco’s Modified Eagle Medium (DMEM) supplemented with 10% FBS; HeLa cells were grown in Minimum Essential Medium (MEM) supplemented with 10% FBS. Cells were maintained at 37°C, 5% CO_2_ in a humidified incubator and subcultured once every 2-3 days or when approaching confluence.

For BRET and flow cytometry experiments, on Day 1, HEK293T or HeLa cells were seeded in 6-well plates at 800K-1M/well or 400K-500K/well, respectively, 6-8 hours prior to transfection. Transfection was performed with TransIT-LT1 for HEK293T and TransIT-X2 for HeLa. Transfection mixtures were prepared continuing a total of 2.5 μg of DNA/well and 7.5 μl of transfection reagent, in 250 μl of Opti-MEM, and incubated for 30 minutes at RT before adding dropwise to respective wells. The DNA ratios used for the transfections for different assays were as follows:

- **CAMYEL assay:** CAMYEL or CAMYEL-V2: 40%, receptor: 20%, Gαi or Gαs: 40%
- **GRK3ct assay:** GRK3ct-RLuc8: 4%, mVenus-Gβ1: 24%, untagged Gγ2: 24%, receptor: 24%, Gαi: 24%
- **G**α**i/G**βγ **assay (three-plasmid):** receptor: 24%, Gαi: 4%, mVenus-Gβ1: 36%, untagged Gγ2: 36%.
- **G**α**i/G**βγ **assay (tricistronic):** receptor: 25%, pcDNA: 25%, pIRES Gβ-2A-cpV-Gγ2 Gαi-Nluc: 50%.
- **G**α**i/G**βγ **assay (three-plasmid) with PTCH1:** receptor: 15%, mPTCH1b-CLIP or pcDNA: 43%, Gαi: 3%, mVenus-Gβ1: 21%, untagged Gγ2: 18%.
- **G**α**i/G**βγ **assay (tricistronic) with PTCH1:** receptor: 15%, mPTCH1b-CLIP or pcDNA: 45%, pIRES Gβ-2A-cpV-Gγ2 Gαi-Nluc: 40%.

24 hours post transfection, on Day 2, cells were lifted from the 6-well plates using Trypsin/EDTA, pelleted by centrifugation, resuspended at a uniform density in phenol-red-free DMEM+10%FBS, and plated in a 96-well plate at 35K-45K cells/well (for HeLa) or 55K-70K cells/well (for HEK293T). The remaining transfected cells were seeded in 6-well plates, to be used for expression control by flow cytometry on the day of the BRET experiment. Replated cells were placed back in the TC incubator and maintained there until the day of the experiment (Day 3). Constitutive activity suppression control wells were pretreated with 100 ng/mL PTX overnight.

### CAMYEL cAMP BRET assay

Cell culture media in 96-well plates with HEK293T or HeLa cells expressing CAMYEL and desired receptors was replaced with a BRET buffer (1X PBS supplemented with 0.1% glucose and 0.05% BSA). The luciferase substrate coelenterazine-h was added to the wells at the final concentration of 5 μM (for HEK293T) or 12-15 μM (for HeLa) with or without 100μM of IBMX. The reagents were mixed by agitating the plate for 15 seconds, and incubated for 3 minutes before initiating repeated readings (every 2.5 min) of light emission at 430-485 nm and 505-590 nm on the TECAN Spark plate reader (Tecan, Switzerland). Basal BRET measurements continued for 6-10 minutes, following the addition of ligands at final concentrations of 50 nM for CXCL12 or SAG21k, 500 nM for IT1t, 500 nM for VUF16840, or 2 μM for Vismodegib and subsequent repeated BRET measurements. Ligand concentrations were consistent across all BRET assays in this study. At ∼8-12 minutes post ligand addition, Forskolin (FSK) was added to all wells at the final concentration of 25 μM and measurements were continued for the additional 20 minutes.

### GRK3ct/G**βγ** association BRET assay

Cell culture media in the 96-well plate with either HEK293T or HeLa cells transiently transfected with GRK3ct-RLuc8, mVenus-Gβ1, and untagged Gγ2 with or without Gαi and receptors of interest, was replaced with the BRET buffer. Immediately before reading, the luciferase substrate coelenterazine-h was added to the wells at the final concentration of 5 μM (for HEK293T) or 12-15 μM (for HeLa). The reagents were mixed by agitating the plate for 15 seconds and incubated for 3 minutes before initiating repeated readings (every 2.5 min) of light emission at 430-485 nm and 505-590 nm on the TECAN Spark plate reader (Tecan, Switzerland). Basal BRET measurements continued for 6-10 minutes, following the addition of agonists at final concentrations of 50 nM (for CXCL12 and SAG21k). At ∼8-12 minutes post agonist addition, inverse agonists were added at the final concentration of 500 nM (for IT1t and VUF16840) or 2 μM (for Vismodegib), and measurements were continued for the additional 35 minutes. Ligand concentrations were consistent across all BRET assays in this study. The collected luminescence emissions were converted into BRET ratio, baseline-corrected by subtracting the average BRET ratio prior to agonist addition, and normalized as described in Figure legends.

### G**α**i/G**βγ** dissociation BRET assay

Cell culture media in the 96-well plate with HEK293T or HeLa cells expressing untagged Gαi, mVenus-Gβ1 and untagged Gγ2 or tricistronic pIRES Gβ-2A-cpV-Gγ2 Gαi-Nluc along with the receptors of interest and/or mPTCH1b-CLIP, was changed to the BRET buffer. Immediately before reading, the luciferase substrate coelenterazine-h was added to the wells at the final concentration of 5 μM (for HEK293T) or 12-15 μM (for HeLa). The reagents were mixed by agitating the plate for 15 seconds and incubated for 3 minutes before initiating repeated readings (every 2.5 min) of light emission at 430-485 nm and 505-590 nm on the TECAN Spark plate reader (Tecan, Switzerland). Basal BRET measurements continued for 6-10 minutes, following the addition of agonists at final concentrations of 50 nM (for CXCL12 and SAG21k). At ∼8-12 minutes post agonist addition, inverse agonists were added at the final concentration of 500 nM (for IT1t and VUF16840) or 2 μM (for Vismodegib), and measurements were continued for the additional 35 minutes. Ligand concentrations were consistent across all BRET assays in this study. The collected luminescence emissions were converted into BRET ratio, baseline-corrected by subtracting the average BRET ratio prior to agonist addition, and normalized as described in Figure legends.

### Flow cytometry

Cells were detached from the 6-or 12-well plates with Accutase and lifted in MEM+10% FBS. All subsequent steps were performed on ice. Cells were washed twice with PBS+0.5% BSA (FACS buffer). For experiments involving antibody-based detection of intracellular epitopes, cells were fixed with PBD+1%PFA for 30 minutes at RT, followed by permeabilization in PBS+0.5%BSA+0.2%Tween-20 for 30 minutes on ice. For permeabilized cells, all subsequent washing and staining steps were also performed in the FACS buffer with 0.2% Tween-20. Samples that required staining (as opposed to those expressing genetically encoded fluorophores like mVenus) were stained with anti-HA-APC (1:50) or anti-Flag-APC (1:50) for 30 minutes on ice, followed by three washes in a relevant FACS buffer with or without Tween-20. Flow cytometry data was collected on Guava benchtop flow cytometer (EMD Millipore). For samples stained with APC-conjugated antibodies, fluorescence was measured in the RED-R channel (ex: 642 nm, em: 695/50 nm). The expression levels of fluorescent BRET acceptors were measured in the GRN-B channel (ex: 488 nm, em: 525/30 nm). Data was analyzed in FlowJo v10.8.0 (FlowJo LLC, Ashland, Oregon).

### Immunofluorescence

HeLa cells were plated on a 12-well plate for 9 hours prior to transfection with Gαi: 20%, mPTCH1b-CLIP or pcDNA: 60%, SMO: 20%. Next day, cells were replated on 12 mm glass coverslips pre-coated with fibronectin (25 μg/mL) for 1 hour at 37 °C, and allowed to attach and grow for 24-48 hours in a 37 °C, 5% CO_2_ incubator. On the day of the experiment, cells on coverslips were rinsed with PBS, fixed in 2% paraformaldehyde/PBS for 30 minutes at RT, washed three times with PBS, and permeabilized/blocked with PBS+1% BSA+0.1% TritonX- 100 for 30 minutes at RT. Next, coverslips were incubated with rabbit anti-SNAP antibodies (1:1000, CAB4255, Invitrogen) diluted in 0.1% TritonX-100/1%BSA/PBS blocking buffer for 1 hour at RT. Coverslips were washed three times with PBS before incubating with secondary antibody AF488-goat anti-rabbit (Thermo A11034, 1:1000) and AF594-conjugated mouse anti-HA (BioLegend 901511, 1:400) diluted in blocking buffer, followed by DAPI for 1 hour at RT. Coverslips were then washed in PBS and affixed to glass slides using ProLong Diamond Antifade mounting solution (P36965, Invitrogen). Images were acquired on a Nikon Eclipse Ti microscope with a 60× Plan Fluor oil immersion objective.

## Abbreviations

GPCR: G protein coupled receptor
GiPCR: Gi protein coupled receptor
GsPCR: Gs protein coupled receptor
GAP: GTPase-Activating Protein.
GDI: Guanine Nucleotide Dissociation Inhibitor
GDP: Guanosine Diphosphate
GTP: Guanosine Triphosphate
FSK: Forskolin
AC: Adenylyl Cyclase
BRET: Bioluminescence Resonance Energy Transfer
cAMP: Cyclic Adenosine Monophosphate
PTX: pertussis toxin
IBMX: 3-isobutyl-1-methylxanthine
CAMYEL: cAMP sensor using YFP-Epac-RLuc
CXCR4: C-X-C chemokine receptor type 4
CXCL12: C-X-C chemokine ligand 12 (same as SDF1)
SDF1: stromal cell derived factor 1 (same as CXCL12)
SMO: Smoothened receptor
ACKR3: Atypical chemokine receptor 3
HA/ha: Hemagglutinin
PTCH1: Patched-1 receptor
GRK: G-protein-coupled receptor kinase
GRK3ct: G-Protein-Coupled Receptor Kinase 3 C-Terminus
RhoA: Ras Homolog Family Member A
HEK293T: Human Embryonic Kidney 293T cells
HeLa: Human epithelial adenocarcinoma cell line
PM: Plasma Membrane
RLuc: Renilla Luciferase
CR: Catalytically Reduced
RR: Regulatory Reduced
EPAC: Exchange Protein Directly Activated

## Data analysis

Data visualization and statistical analyses were performed in GraphPad PRISM 10.0.2. Statistical tests are described individually in the figure legends.

## Acknowledgements

The authors are grateful to Paul Insel and Pradipta Ghosh, UC San Diego, Ben Myers, Univ. of Utah, Mikel Garcia-Marcos, Boston Univ., Michel Bouvier, Univ. of Montreal, Nevin Lambert, Augusta Univ., Tohoku Univ., and Daniel Legler, Univ. of Konstanz, for generous gifts of receptor and reporter constructs. We thank Beatrice Acot, Christopher Schaefer, and other members of the Kufareva and Handel labs for valuable discussions. This work was supported by NIH grants R21 AI149369, R21 AI156662 (to I.K.) and R01 AI161880, R01 GM136202 (to I.K. and T.M.H.).

## Author contributions

I.K. designed and supervised the research. K.C., J.R., E.K., and S.E. collected data. K.C., J.R., E.K., and S.R. performed molecular cloning. S.E., K.C., and I.K. analyzed the data. S.E. created figures. S.E., I.K., and T.M.H. wrote the paper, with input from all co-authors. All authors approved the final version of the manuscript.

## Supplemental Figures and Legends

**Supplementary Figure 1.**
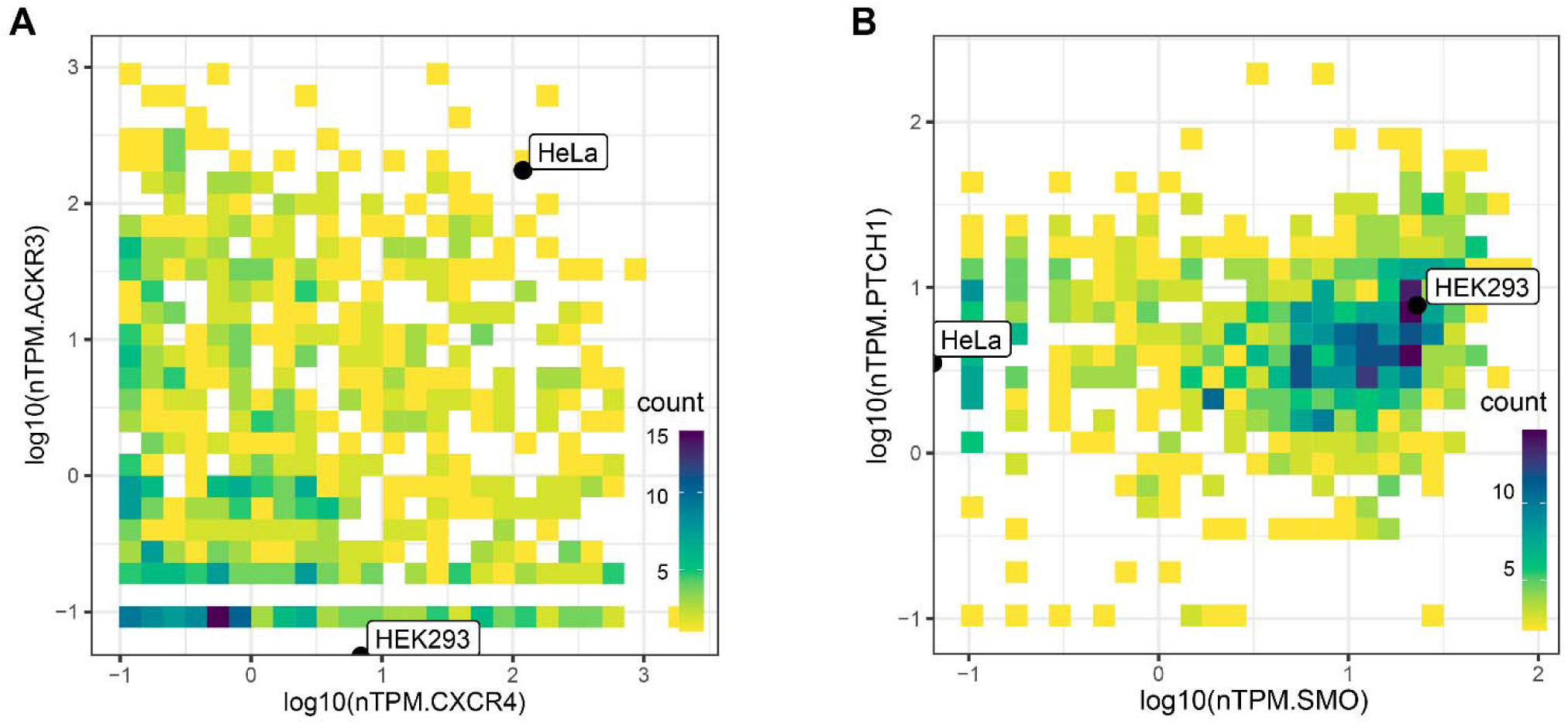
Endogenous co-expression of receptors in this study in various cell lines. (**A, B**) Normalized mRNA expression (in Transcripts Per Million, nTPM) of CXCR4 vs ACKR3 (**A**) and SMO vs PTCH1 (**B**) in a panel of 1055 cell lines from the Human Protein Atlas [86]. HeLa and HEK293T cells used in the study are labeled.

**Supplementary Figure 2.**
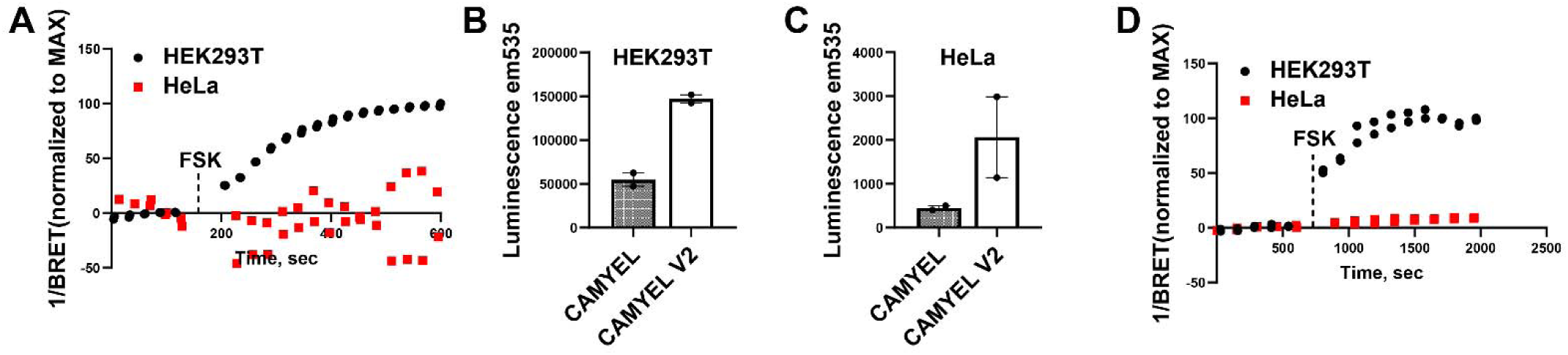
CAMYEL vs CAMYEL-V2 biosensor. (**A**) Mapping the time course of changes in CAMYEL 1/BRET ratios in HEK293T or HeLa cells upon the addition of Forskolin. (**B, C**) Absolute emission in the 505-590 nm range for the CAMYEL vs CAMYEL-V2 biosensor in HEK293T (**B**) and HeLa (**C**) cells, measured at 5-8 min post CTZ-h addition. (**D**) Time course of changes in CAMYEL-V2 1/BRET ratios in HEK293T or HeLa cells upon the addition of Forskolin.

**Supplementary Figure 3.**
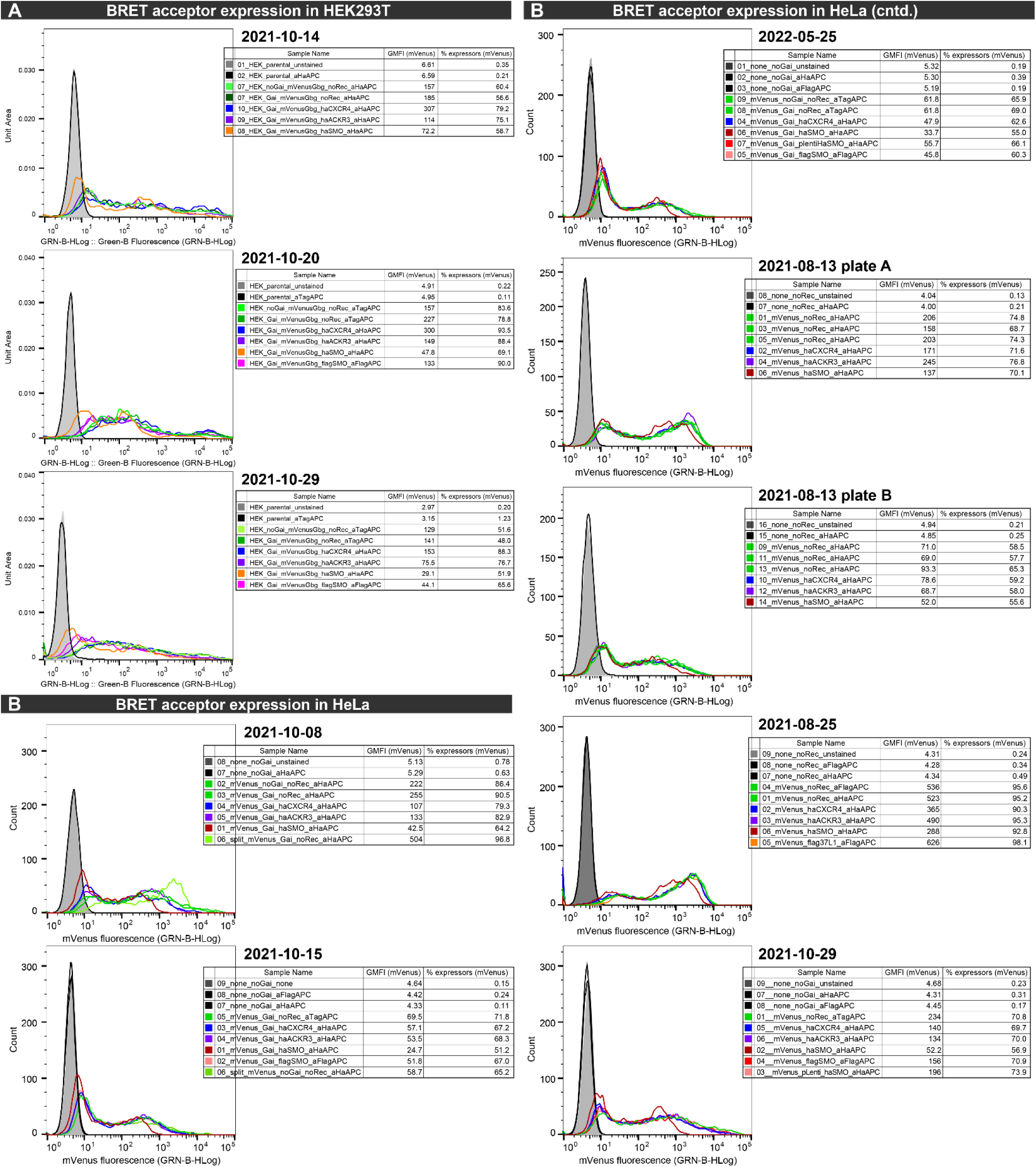
Quantification of BRET acceptor expression by flow cytometry. Flow cytometry assessment of BRET acceptor expression in samples co-transfected or not with the indicated receptors, in individual biological replicates of BRET experiments presented in the paper for HEK293T cells (**A**) and HeLa cells (**B**).

**Supplementary Figure 4.**
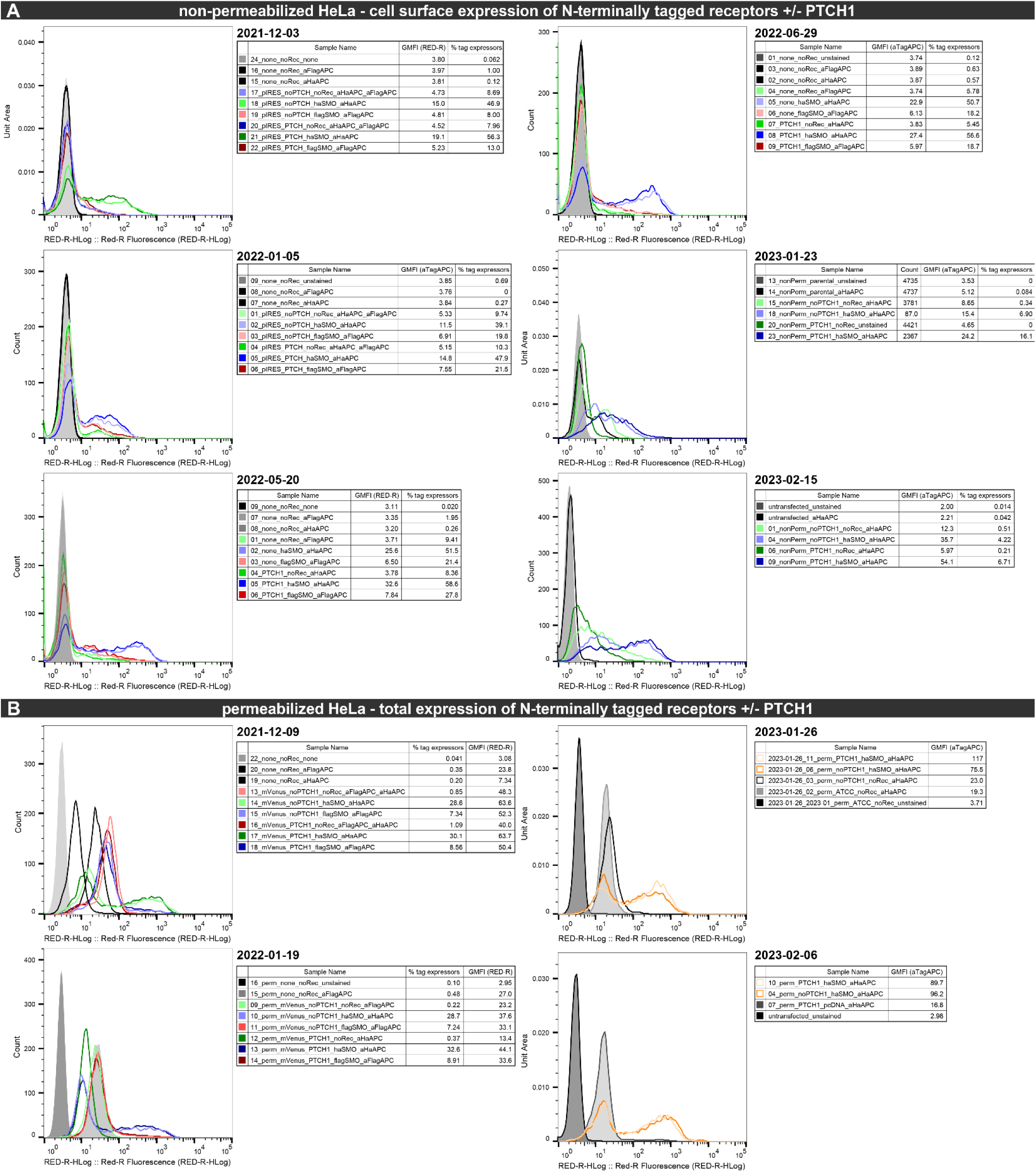
Quantification of SMO expression in the presence or absence of PTCH1 by flow cytometry. Flow cytometry quantification of surface (**A**) or total (**B**) expression of indicated SMO constructs in HeLa cells in independent biological experiments. N-terminally HA- or Flag-tagged receptors were detected using an APC-conjugated anti-HA antibody.

**Supplementary Figure 5.**
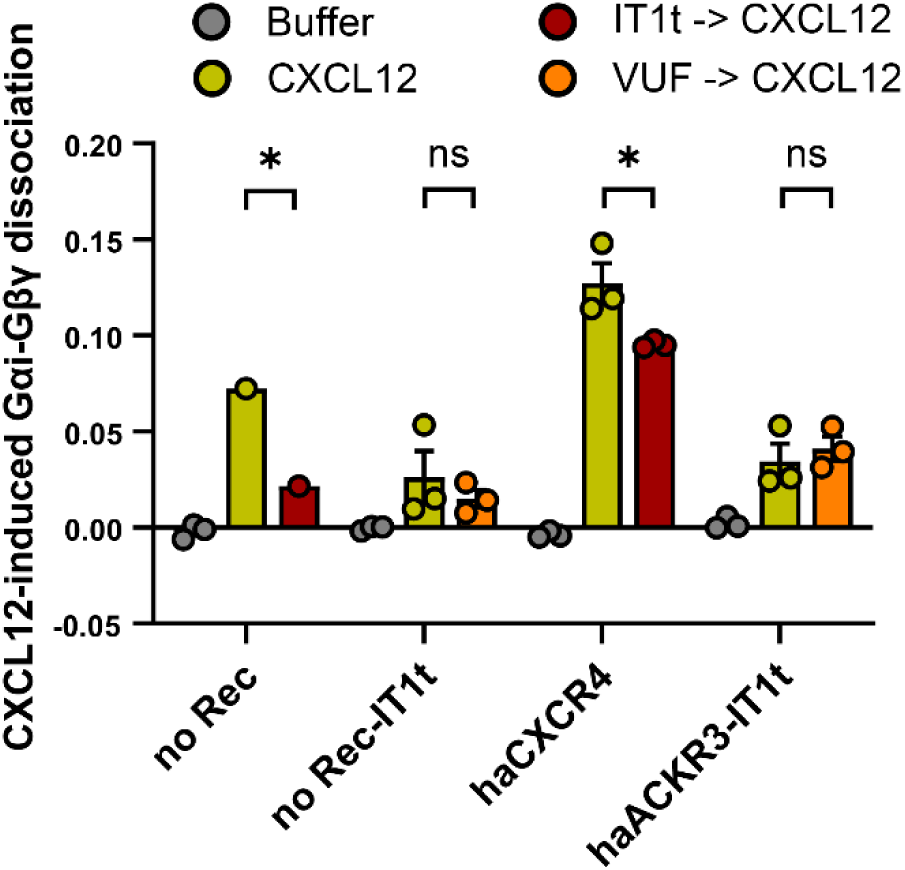
CXCL12-induced changes in Gαi-Gβγ dissociation. CXCL12-induced changes in Gαi-Gβγ dissociation in HEK293T cells transfected with the indicated receptors pre-treated or not with 500nM IT1t (a CXCR4 antagonist) or VUF16840 (an ACKR3 antagonist).

